# Cryo-EM structures of pentameric autoinducer-2 exporter from *E. coli* reveal its transport mechanism

**DOI:** 10.1101/2021.10.20.465058

**Authors:** Radhika Khera, Ahmad Reza Mehdipour, Jani R Bolla, Joerg Kahnt, Sonja Welsch, Ulrich Ermler, Cornelia Muenke, Carol V Robinson, Gerhard Hummer, Hao Xie, Hartmut Michel

**Author notes:** Address correspondence to Hao Xie, or Hartmut Michel.

## Abstract

Bacteria utilize small extracellular molecules to communicate in order to collectively coordinate their behaviors in response to the population density. Autoinducer-2 (AI-2), a universal molecule for both intra- and inter-species communication, is involved in the regulation of biofilm formation, virulence, motility, chemotaxis and antibiotic resistance. While many studies have been devoted to understanding the biosynthesis and sensing of AI-2, very little information is available on its export. The protein TqsA from *E. coli*, which belongs to a large underexplored membrane transporter family, the AI-2 exporter superfamily, has been shown to export AI-2. Here, we report the cryogenic electron microscopic structures of two AI-2 exporters (TqsA and YdiK) from *E. coli* at 3.35 Å and 2.80 Å resolutions, respectively. Our structures suggest that the AI-2 exporter exists as a homo-pentameric complex. In silico molecular docking and native mass spectrometry experiments were employed to demonstrate the interaction between AI-2 and TqsA, and the results highlight the functional importance of two helical hairpins in substrate binding. We propose that each monomer works as an independent functional unit utilizing an elevator-type transport mechanism. This study emphasizes the structural distinctiveness of this family of pentameric transporters and provides fundamental insights for the ensuing studies.

## Introduction

Quorum sensing (QS) is a cell-to-cell communication mechanism that confers bacteria with the ability to sense and decipher their surroundings in terms of not only their own cell density, but also other species cohabitating with them^1,2^. This phenomenon was first described for the marine organism *Vibrio fischeri* that emits bioluminescence in response to a high population density^3^. Later, research revealed that the role of QS extends beyond bioluminescence to many other density dependent cellular processes like the formation of biofilms, production of virulence factors and antibiotics, sporulation, conjugation and chemotaxis^1,4–8^. QS specifically modulates those cellular functions and processes, the effect of which will be maximum when all the cells act in unison.

In QS, intercellular communication is mediated by small chemical molecules known as autoinducers^9^. They are synthesized intracellularly and could be passively or actively released into the extracellular milieu. The extracellular concentration of the autoinducer increases as a function of cell number. On reaching a critical concentration threshold, the autoinducers are perceived by their cognate membrane receptors. The bacteria eventually respond to this environmental stimulus by activating signal transduction cascades that ultimately regulate the expression of certain genes. So far, a number of different autoinducers have been identified in various bacteria and they can be classified into three main categories. Acyl-homoserine lactones (AHLs, autoinducer-1) are frequently used by Gram-negative bacteria, while autoinducing peptides (AIPs) are the key autoinducers in Gram-positive bacteria^10^. These signaling molecules display a large structural diversity and are often specific to each bacterial species. There is a third category of autoinducers, termed Autoinducer-2 (AI-2), which can be produced and sensed by a majority of bacterial and eukaryotic species and hence is considered as a universal signal molecule, promoting both intra- and inter-species as well as interkingdom communication^11–13^.

Bacterial AI-2 molecules are small cyclic furanone compounds that are converted from their precursor, 4,5-dihydroxy-2,3-pentanedione (DPD)^14^. DPD is synthesized in three enzymatic steps from *S*-adenosyl methionine (SAM), a metabolite involved in many metabolic pathways, including transmethylation, transsulfuration, and aminopropylation. In the first step, SAM is converted to *S*-adenosyl homocysteine (SAH) by methyltransferases. Subsequently, the SAH nucleosidase hydrolyzes SAH to form *S*-ribosylhomocysteine and adenine. Finally, SRH is converted to DPD and homocysteine by the S-ribosylhomocysteinase (LuxS) enzyme^11^. DPD is quite unstable and can spontaneously cyclize to form different cyclic furanone compounds. Among them, a furanosyl borate diester, called *S*-2-methyl-2,3,3,4-tetrahydroxy tetrahydrofuran-borate (*S*-THMF-borate), was reported to be the AI-2 signal molecule responsible for QS in *Vibrio* species, whereas a non-borated enantiomer of DPD, *R-2*-methyl-2,3,3,4-tetrahydroxy tetrahydrofuran (*R*-THMF), represents the AI-2 signal in a range of other bacteria, including *Salmonella* Typhimurium and *Escherichia coli*^15,16^. AI-2 is considered to be a universal communication signal because LuxS, the key enzyme in AI-2 biosynthesis, was found to be widespread in both Gram-negative and Gram-positive bacteria^17^. This enzyme is also a part of the essential activated methyl cycle, raising skepticism about the idea that AI-2 is just a metabolic byproduct. Subsequent studies revealed that AI-2 is involved in regulation of biofilm formation, pathogenesis, chemotaxis and antibiotic resistance^11,18–22^.

AI-2 or DPD is a hydrophilic compound and is considered to be membrane impermeable^23^. Cell signaling via AI-2 involves both secretion and sensing that require the transmission of signals across the membrane in both directions, as illustrated in Fig. 1. In enteric bacteria, the active import of AI-2 into the cell is carried out by an ABC-type transporter, whose encoding genes are part of the *lsr* operon consisting of eight genes, *lsrKRACDBFG*, that encodes proteins involved in uptake, phosphorylation and further processing of AI-2^14^. However, to the best of our knowledge, the mechanism of export of AI-2 still remains undetermined. The first protein suggested to be an exporter for AI-2 was the YdgG protein (later renamed TqsA for <u>t</u>ransporter of <u>q</u>uorum-sensing <u>s</u>ignal <u>A</u>I-2) from *E. coli*^24^ Previous studies showed that a deletion of *tqsA* leads to increased biomass production, motility, biofilm formation, intracellular concentrations of AI-2 and surged *lsr* operon expression^24^. TqsA is a 38 kDa transmembrane protein, which belongs to a large and underexplored family of putative membrane transporters, presently known as the AI-2 exporter superfamily. In *E. coli*, three more proteins (YhhT, YdiK and PerM) were assigned to this superfamily^25^. The expression of two of them (PerM and YdiK) was shown to be regulated by the PurR transcription factor that regulates purine metabolism and transport^26^. The regulation of PurR on the expression of AI-2 exporters at the cellular level is surely noteworthy. The rationale behind this could either be the production of adenine as a by-product during the AI-2 biosynthesis or could indicate the existence of an additional set of substrates (purines) for this exporter family^25^.

**Fig. 1.**
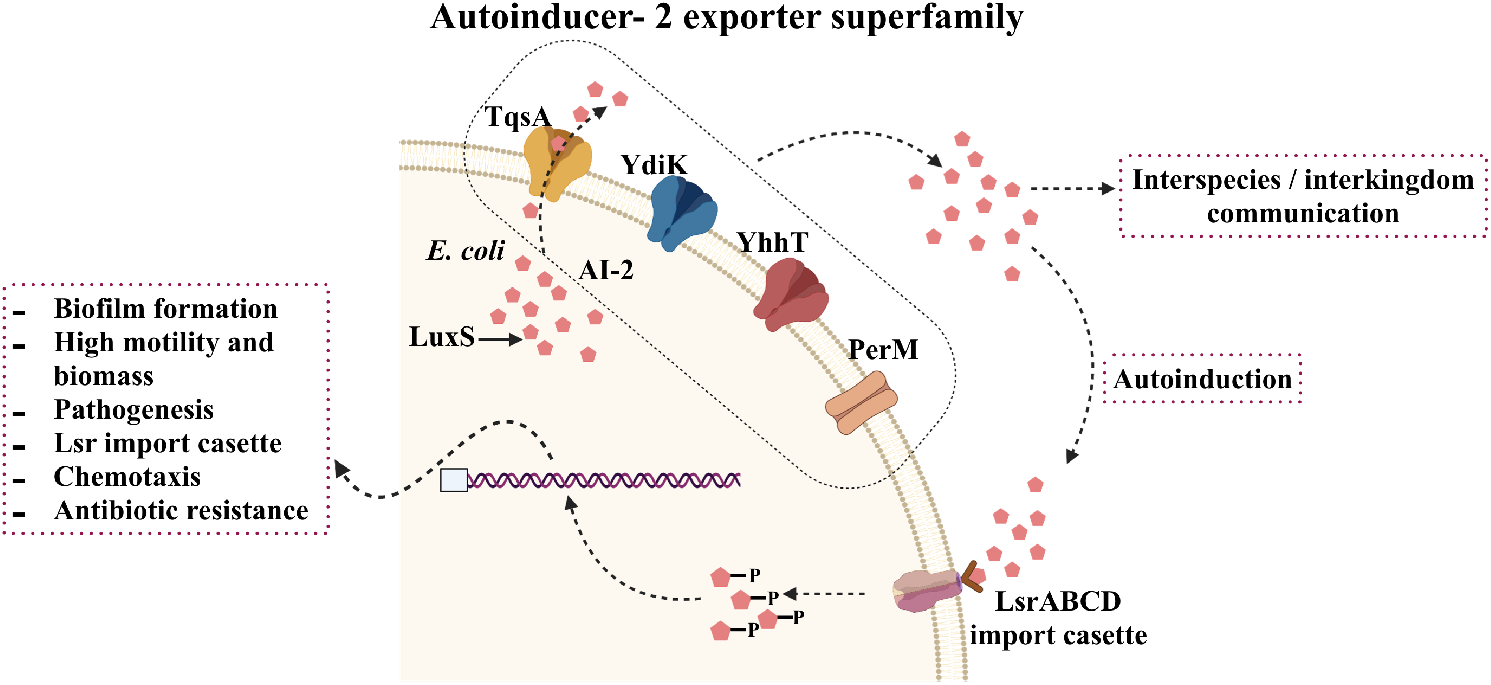
Schematic overview of AI-2 mediated quorum sensing in *E. coli*. Autoinducer-2 is quorum sensing signal that mediates interspecies and interkingdom cell-to-cell communication. The four predicted AI-2 exporters from *E. coli* are shown embedded in the inner membrane in different shades (TqsA in yellow, YdiK in blue, YhhT in red and PerM in brown), AI-2 is represented as small red pentagonal moieties. LuxS produces DPD, the precursor of AI-2. The ABC transporter complex LsrABCD is involved in the active import of AI-2. Phosphorylated AI-2 inactivates the transcriptional repressor and thereby regulates the expression of quorum sensing-related genes.

In this study, all four AI-2 exporters from *E. coli* (TqsA, YhhT, YdiK and PerM) were purified and studied in an attempt to characterize them both biochemically and structurally. We determined the cryogenic electron microscopy (cryo-EM) structures of TqsA and YdiK at 3.35 Å and 2.80 Å resolution, respectively. Our results show that these AI-2 exporters are homo-pentameric complexes, and structural analyses in combination with molecular docking studies reveal the importance of two helix-turn-helix motifs for substrate binding. This study provides major insights into this novel and unexplored superfamily of membrane transporters and will serve as the basis for future studies aimed at understanding the mechanisms of AI-2 secretion.

## Results

### AI-2 exporters from *E. coli*

The proteins of the AI-2 exporter superfamily are approximately 40 kDa in size with a few exceptions^25^. The results of multiple sequence alignment of the four predicted AI-2 exporters from *E. coli* (TqsA, YdiK, YhhT and PerM) showed that each protein is distantly related to each other with a relatively low overall sequence similarity (Fig. S1 f). The highest sequence identity of 46% was observed between TqsA and YhhT, indicating a closer phylogenetic relationship. No clear phylogenetic correlation can be drawn between TqsA, YdiK and PerM since pairwise sequence identities fall below ≈24%, which may suggest different cellular functions or substrate specificities of the proteins.

Recently, a member of the AI-2 exporter superfamily from *Halobacillus andaensis* (Upf0118) was found to function as a Na^+^/H^+^ antiporter, which represents an independent subgroup in the AI-2 exporter superfamily^27,28^. A comparison between TqsA and Upf0118 revealed a low sequence similarity of 24%. Nevertheless, we investigated if the four selected AI-2 exporters from *E. coli* can function in a similar manner as Upf0118. Results of our growth complementation assay showed that the *E. coli* Na^+^-sensitive strain (Knabc strain) expressing the AI-2 exporters was unable to grow in the presence of sodium ions, which suggests that they do not act as Na^+^/H^+^ antiporters (Fig. S12 a, b). In addition, the assay of antiport activity using everted vesicles further confirmed that TqsA does not transport Na^+^/Li^+^ in exchange for H^+^ (Fig. S12 c, d).

### Purification and characterization of four AI-2 exporters

In this work, four AI-2 exporters were produced in *E. coli* Top10 cells. Initially, the detergent GDN was used for the purification. All four proteins could be purified to homogeneity at yields of sub-milligram proteins per liter of culture using a combination of affinity and size-exclusion chromatography (Fig. S1 a-d). Gel filtration profiles showed that three AI-2 exporters (TqsA, YdiK and YhhT) eluted as single sharp peaks at similar retention volumes of ~15 ml corresponding to a size of about 400 kDa (Fig. S1 a-c), while PerM was recovered later from the column as a broader peak at 16.2 ml (Fig. S1 d), suggesting the presence of different oligomeric species with the majority possessing apparently smaller hydrodynamic radii.

All purified proteins were analyzed on BN-PAGE gels to investigate their oligomeric state (Fig. S1 a-d). Three proteins (TqsA: 37.5 kDa; YdiK: 39.8 kDa; YhhT: 38.5 kDa) migrated as single dominant bands with apparent molecular masses of about 320 kDa (Fig S1 a-c), which corresponds to around 180 kDa using a correction factor of 1.8 to account for the mass contribution of bound Coomassie dye and detergent molecules^29^. The deduced mass suggests that all three proteins may exist as homo-oligomeric complexes consisting of four or five protomers. By comparison, a single dominant band migrating at approximately 155 kDa was observed for PerM (39.2 kDa), yielding a mass of ~86 kDa after correction, which may indicate the presence of a dimer.

To investigate the influence of detergents on the oligomerization of AI-2 proteins, several non-ionic detergents (α-DDM, β-DDM, DM and LMNG) were also utilized for the purification. After solubilization with β-DDM, detergent exchange was directly performed on the affinity chromatography column. The oligomeric states of the purified proteins were further assessed by size exclusion chromatography and BN-PAGE analysis. All proteins displayed similar behavior to that described in GDN (Fig. S1 a-d), indicating that the assembly of higher oligomeric complex was not affected by the choice of detergents.

### Single-particle cryo-EM analyses of AI-2 exporters

As already mentioned, monomers of AI-2 exporter proteins are relatively small, and the size is close to the theoretical lower protein size limit for the visualization in cryo-EM^30^. Nevertheless, the existence of higher oligomeric states of AI-2 exporters allowed us to successfully execute electron microscopic high-resolution structure determinations without the need for additional fusion proteins or binding proteins. For the cryo-EM experiments, we attempted protein preparation in the two commonly used detergents LMNG and GDN. After data processing, we compared different datasets, and we found that GDN was more suitable for the AI-2 exporters than LMNG, yielding a better quality of reconstructed 3D density maps. A total of 5,452 and 7,372 movie stacks were collected for GDN-purified TqsA and YdiK, respectively (Fig. S2 a). The two-dimensional (2D) class averages showed that TqsA adopts a characteristic pentameric structure resembling the shape of a “flower” with five petals when viewed from the top and display a bowl-shaped architecture equipped with a central cavity when viewed from the side (Fig. S2 b). YdiK also forms a similar pentameric arrangement, however, the central cavity is not that pronounced in the 2D class averages (Fig. S2 b). The final three-dimensional (3D) reconstructions yielded maps with 3.3 Å and 2.8 Å resolution for the overall structures of TqsA and YdiK, respectively (Fig. 2 a, c). In both EM maps, the bottom and the interior of the bowl are the best resolved portions of the structure (~3.0 Å for TqsA and ~2.7 Å for YdiK), while the rim of the bowl is not that well resolved, especially in the EM map of YdiK, probably due to the high flexibility of this region (Fig. S2 c, f). Finally, we built the atomic models de novo for both AI-2 exporters (Fig. 2 b, d). Our final model for each TqsA monomer contains 340 out of a total of 344 residues, spanning Ala2 to Leu341 (Fig. S7 a). For YdiK, some portions of the protein could not be modelled due to lack of density, and the final model includes three polypeptides, Pro7-Ile80, Phe163-Gly230 and Thr279-Gln354, comprising ~ 59 % of the total amino acid residues (Fig. S7 b).

**Fig. 2.**
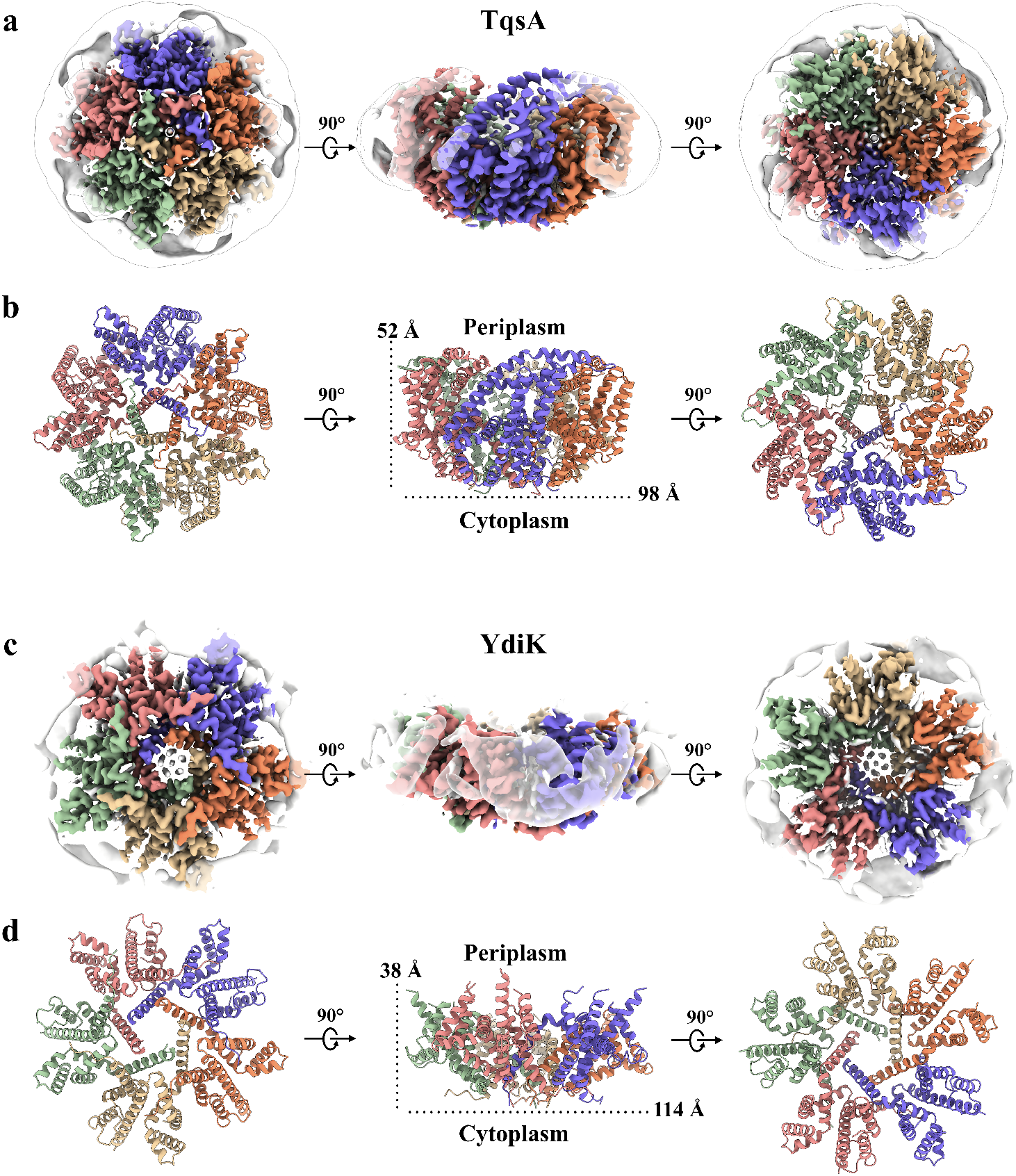
Cryo-EM structures of the pentameric complex of the two AI-2 exporters. **a:** Cytoplasmic (left), side (middle) and periplasmic (right) view of a cryo-EM 3D reconstruction map of TqsA at 3.35 Å resolution colored by protomers. The detergent belt surrounding the transmembrane helix region of the protein is shown. **b:** Corresponding structural model of the TqsA pentamer. Protomers are colored as in **a**. **c:** The 2.80 Å cryo-EM density map of YdiK showing the pentameric organization of YdiK. Each protomer is presented in a different color. **d:** Corresponding structural model of YdiK pentamer. Protomers are colored as in **c**.

In addition, a small set of 989 movies were collected for the GDN-purified YhhT (Fig. S2 a). This dataset exhibited a pronounced preferred orientation, presenting mainly side-views of the particles in 2D class averages. Due to the absence of top-views, direct visualization of the oligomeric state of YhhT was not possible (Fig. S2 b). Nevertheless, YhhT exhibits very similar side-views as observed for TqsA and YdiK. The particle size of all three proteins has similar dimensions of 135-145 Å in diameter, thereby providing sufficient evidence for a pentameric nature of YhhT (Fig. S2 b).

### Pentameric architecture and oligomeric interface

The cryo-EM structures of TqsA and YdiK revealed that both AI-2 exporters form a pentamer with the five-fold non-crystallographic symmetry axis perpendicular to the membrane (Fig. 2 b, d). The structure of the pentameric complex features a bowl-shaped architecture with a concave aqueous basin facing the periplasmic space and a tightened base located in the cytoplasm. The TqsA pentamers are about 98 Å in diameter and 52 Å in height (Fig. 2 b). On the periplasmic side, the rim of the bowl is shaped like a pentagon with each side approximately 39 Å in length. The basin has a depth of around 30 Å and penetrates the membrane bilayer from the periplasm about two-thirds. The concave surface is mainly hydrophilic which may allow the bulk aqueous solution to access the membrane interior to a considerable depth (Fig. S9 a). Between the two TqsA protomers within the pentamer, the interface has an area of 1717 Å^2^ per monomer and is mainly formed by hydrophobic interactions of residues present in the N-terminal regions from the two adjacent monomers (Fig. S6 d). In addition, the interface is further strengthened by a few H-bonds, *e.g*., between Ser145 (protomer A) and Tyr127 (protomer B) on the periplasmic side as well as between Lys171 (protomer A) and Leu337/Ser339 (protomer B) on the cytoplasmic side.

The overall structure of the YdiK pentamer, which is 114 Å in diameter, is similar to that of the TqsA protein (Fig. 2 d). However, the height of the pentameric YdiK and the depth of its basin cannot be accurately estimated since the resolved structure was only partially modelled and the majority of the missing density lines the periplasmic basin. Nevertheless, one interesting structural feature can be recognized at the cytoplasmic bottom of the pentamer in both proteins. There is a pentagonal central pore connecting the cytoplasm with the periplasmic aqueous basin (Fig. 4 a, b). The pentagonal opening at the cytoplasmic side has a width of 17 Å with each side of the pentagon of 11 Å for TqsA and a width of 20 Å with each side of the pentagon of 11.6 Å for YdiK (Fig. 4 a, b). The pore spans vertically into the membrane to a depth of 22 Å for TqsA and 17 Å for YdiK, and subsequently it opens up in the basin facing the periplasm. The overall shape of this pore is dissimilar in both AI-2 exporters. It resembles a cylinder in the case of TqsA, while it appears more like a funnel opening up in the periplasmic basin in YdiK (Fig. 4 a, b). The pore widens up a bit in both proteins while extending perpendicularly in the membrane, but it widens significantly more in terms of area in the case of YdiK (around five times) in comparison to TqsA which widens to almost twice the size of its cytoplasmic opening. The pore in both proteins is constituted by Helix1 (H1) and this helix from each protomer entangles against the neighboring one. This helix mostly comprises hydrophobic residues resulting in an overall hydrophobic environment in the central cavity of the pore (Fig. 4 c, d).

**Fig. 3.**
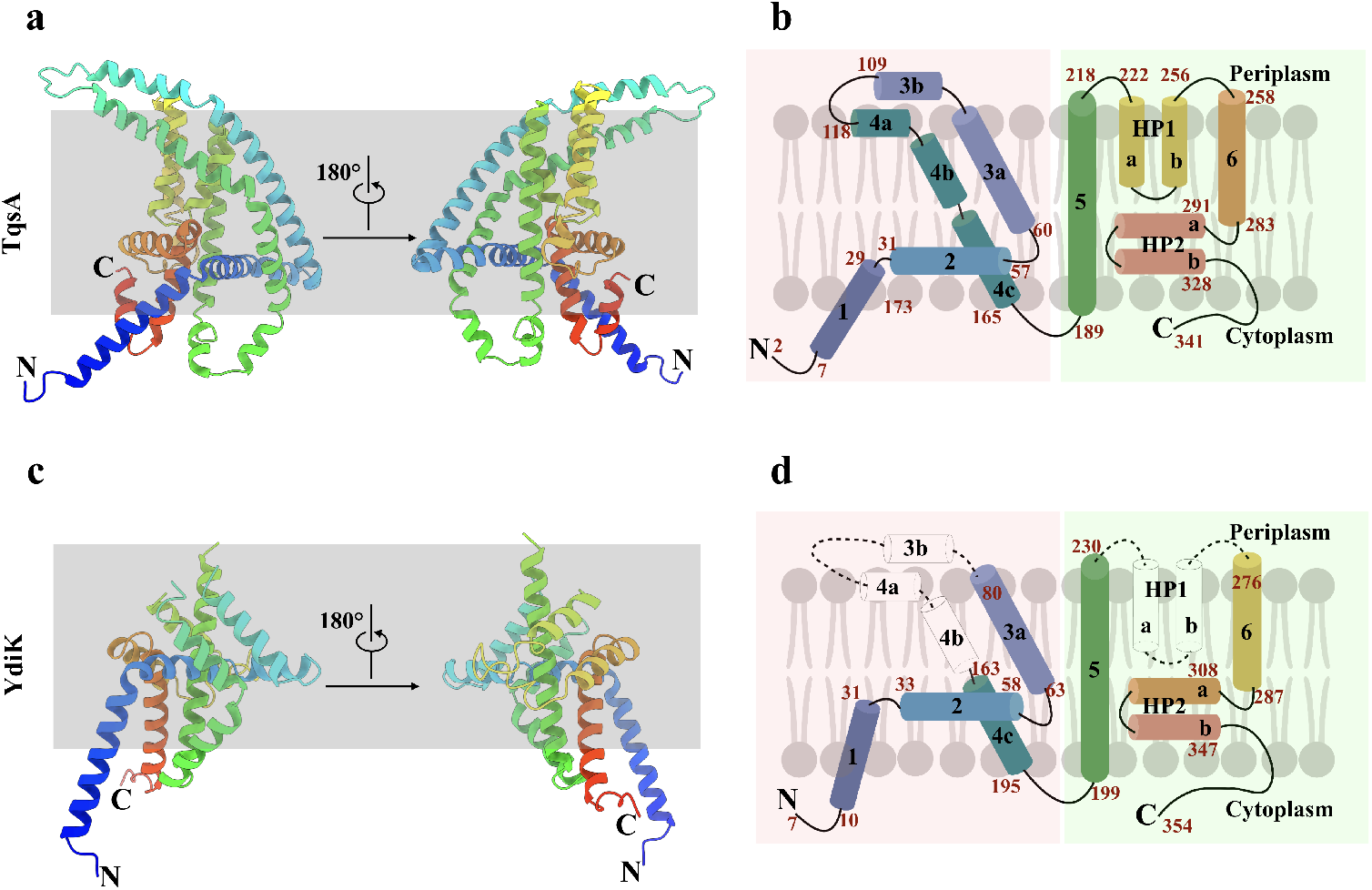
Atomic structures of the two monomeric AI-2 exporters. **a:** Ribbon representation of the TqsA monomer, viewed from the membrane using a rainbow color gradient from the N- (blue) to C-terminus (red). **b:** Schematic representation of the transmembrane topology of TqsA. α-helices are shown as cylinders, sequence numbers of amino acids of segments are marked in red. The N- and C-terminal domains are shaded in red and green, respectively. **c, d:** Structure of the YdiK protomer as shown for TqsA in figures **a** and **b**. The hydrophilic part of TM3 and TM4 as well as HP1 are not included in the atomic model due to the weak density for this part in the cryo-EM map. The missing loops are represented as dashed connecting lines and missing helices are shown as white cylinder in the topology map.

**Fig. 4.**
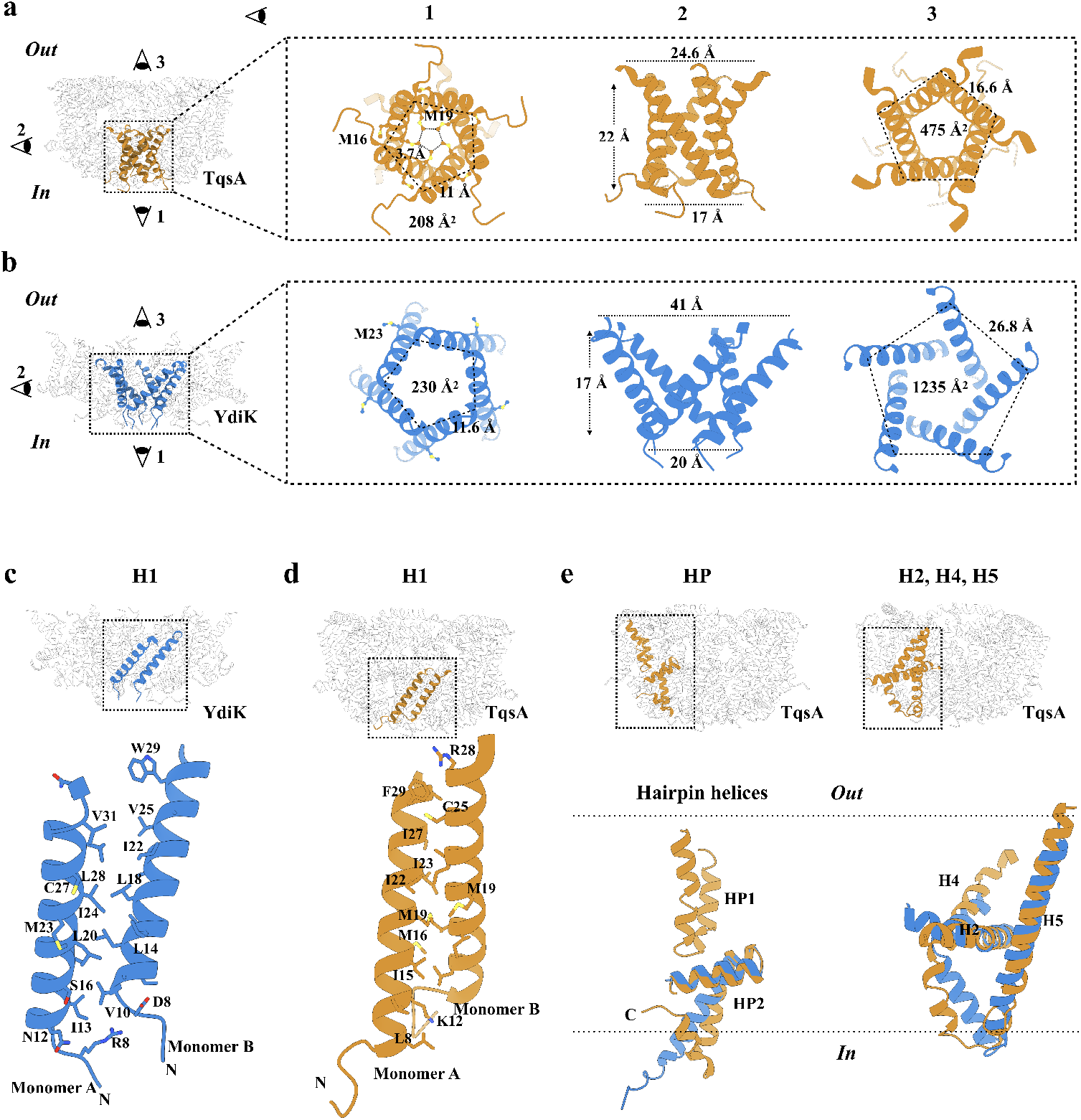
Structural features of the AI-2 exporters TqsA and YdiK. **a, b:** The figures focus on the cytoplasmic central pore of the proteins TqsA (**a**, brown) and YdiK (**b**, blue). Images on the left in both figures show the overall positioning of the pore in the pentameric complexes. The illustrations on the right represent distinct views of the pore from three different directions. **c, d:** Close-views showing the hydrophobic interactions between two adjacent H1 helices, which are vital for maintaining the pentameric architecture of the AI-2 exporters. **e:** Structural superimposition of TqsA and YdiK in specific regions of the proteins in order to highlight their structural deviations. Helical hairpins are shown on the left while H2, H4 and H5 on the right.

We could observe long non-protein densities in the central cytoplasmic pore of YdiK and around it in the case of TqsA, suggesting the presence of either lipids or detergent molecules. In the central pore of TqsA, there is a methionine residue (Met19^TqsA^) facing the interior of the pore from each protomer as shown in Fig. 4 a. The distance between the adjacent residues is ~3.7 Å and the arrangement of five methionine side chains appears to be like a hydrophobic filter in the cavity. Methionine residues, Met16^TqsA^ and Met23^YdiK^, are also present in the central pore region, but they face the exterior of the pore unlike Met19^TqsA^ (Fig. 4 a, b). The presence of this methionine hydrophobic filter in TqsA and lipids/detergents in the case of YdiK, appears to prevent any possible leakage of ions or water molecules across the membrane through such wide openings in both proteins.

### Overall structure of TqsA and YdiK protomer

The TqsA protomers are mainly composed of α-helical segments with both N- and C-terminus exposed to the cytoplasmic side of the membrane (Fig. 3 a). Viewed parallel to the membrane plane, the protein has the shape of a “half-moon”. Each protomer can be clearly divided into an N-terminal domain formed by four α-helices (H1-4) and a C-terminal domain consisting of two transmembrane α-helices (TMH5 and 6) as well as two helical hairpin motifs (HP1 and HP2) as shown in Fig. 3 b.

The fold of the N-terminal domain is less compact than that of the C-terminal half of TqsA and displays several unusual structural features, which are not commonly observed in other membrane transporters. H1 (Thr7-Phe29) is inclined to the membrane normal by about 40° and penetrates only partially into the lipid monolayer (Fig. 3 a). H1 from all five protomers entwine obliquely around the five-fold axis, forming the major contact point on the cytoplasmic face of the pentamer (Fig. 4 a). This N-terminal helix unravels at Ala30, followed by the membrane-embedded horizontal helix H2. The 42 Å-long H2 (Ala31-Trp57) is oriented nearly parallel to the membrane plane and is localized predominantly in the inner monolayer. This horizontal helix H2 forms an angle of about 105° with H1 and an angle of 60° with H3. The two long helices H3 (Arg61-Glu109) and H4 (Asp118-Glu164), which form an antiparallel helical bundle, constitute another unusual structural element in the N-terminal domain of TqsA. Each helix is broken into segments that result in a highly curved overall shape (Fig. 3 a, b). Both helices can be divided into two approximately equal parts with the hydrophobic portion spanning the membrane and a hydrophilic region exposed to the periplasm. Within the membrane, this two-helix bundle is tilted from the membrane normal by approximately 45° (Fig. 3 a, b). In the TM domain of H3 and H4, mainly van der Waals interactions are responsible for the helix-helix interactions. Interestingly, several leucine residues are spaced regularly at three or four amino acid intervals, and this leucine zipper-like dimerization motif may further stabilize the bent conformation of this two-helix bundle (Fig. S6 b). At the membrane-water interface, H3 and H4 are tilted significantly away from the membrane normal and finally end almost parallel to the membrane surface at the periplasmic side. The connecting loop between H3 and H4 forms one of the major contact points between two adjacent protomers on the boundary of the periplasmic basin (Fig. S6 d). The density of this loop was quite feeble, suggesting the flexibility of this region (Fig. S7 a).

The C-terminal domain of TqsA is compactly assembled into a cylinder-shaped unit. TMH5 (Ala189-Leu218) is oriented perpendicular to the membrane plane. The loop connecting H4 and TMH5 crosses over H2 and is also one of the flexible loops in the structure (Fig. 4 e). TMH5 together with H2 form an uncommon cross-shaped structure, lining the interface between the N- and C-halves and this feature may provide an anchor for possible conformational rearrangements during the transport cycle (Fig. S6 a). Following H5, two helical hairpins HP1 (Phe222-Asn256) and HP2 (Ser291-Thr328) constitute a functionally critical element in the C-half of TqsA (Fig. 3 a, b). Both HP1 and HP2 are helix-turn-helix supersecondary structures mainly composed of hydrophobic residues. HP1 begins on the periplasmic side of the membrane and penetrates vertically about halfway across the membrane. This helical hairpin is sandwiched between H3 and H4 from the N-half and TMH5 and TMH6 from the C-half of TqsA. Both HP1 and HP2 are connected by the 36 Å long helix TMH6 (Phe258-Ile283) followed by a 7 residues loop (Met284-Leu290). Unlike HP1, HP2 is oriented nearly parallel to the membrane plane, with its turn region exposed to the periplasmic basin. Both helical hairpins form an L-shaped structure with the tips pointing in different directions (Fig. 4 e). In HP2, the second helix is highly bent, pointing to the cytoplasmic side. HP2 is followed by the C-terminal region of around 13 residues returning towards the membrane yet exposed in the cytoplasm (Fig. 3 a, b).

Similar to TqsA, each YdiK protomer comprises six α-helical segments and two helical hairpins (Fig. 3 c, d). Due to the lack of density, some portions of the protein could not be modelled, including the hydrophilic region of H3 and H4 and the first helical hairpin (Fig. 3 c, d and Fig. S7 b). Nevertheless, YdiK shows similar helical arrangement to that observed in TqsK (Fig. 4 e). All uncommon structural features that we observed in the TqsA structure, including the membrane-embedded horizontal helix H2, the cross-shaped arrangement between H2 and H5 as well as the placement of HP2, can also be discerned in YdiK (Fig. 4 e).

### Interaction with AI-2

We investigated the interaction between TqsA and its putative substrate AI-2 (*R*-THMF) with *in silico* substrate docking. Our docking results show that the AI-2 compound fits well into the cavity formed by HP1, H5 and H6 in the protomer (Fig. 5 a and Fig. S 9 c). In this binding mode, the AI-2 molecule interacts with the surrounding residues, including Tyr196, Lys200, Pro237, Asn238 and Glu280. Asn238 and the strictly conserved Pro237 are located at the tip of HP1 and they are within the hydrogenbond distance to AI-2 (Fig. 5 a). On the other side of the binding site, the substrate interacts with Glu280, which is semi-conserved among the members of the family (Fig. 5 a, b).

**Fig. 5.**
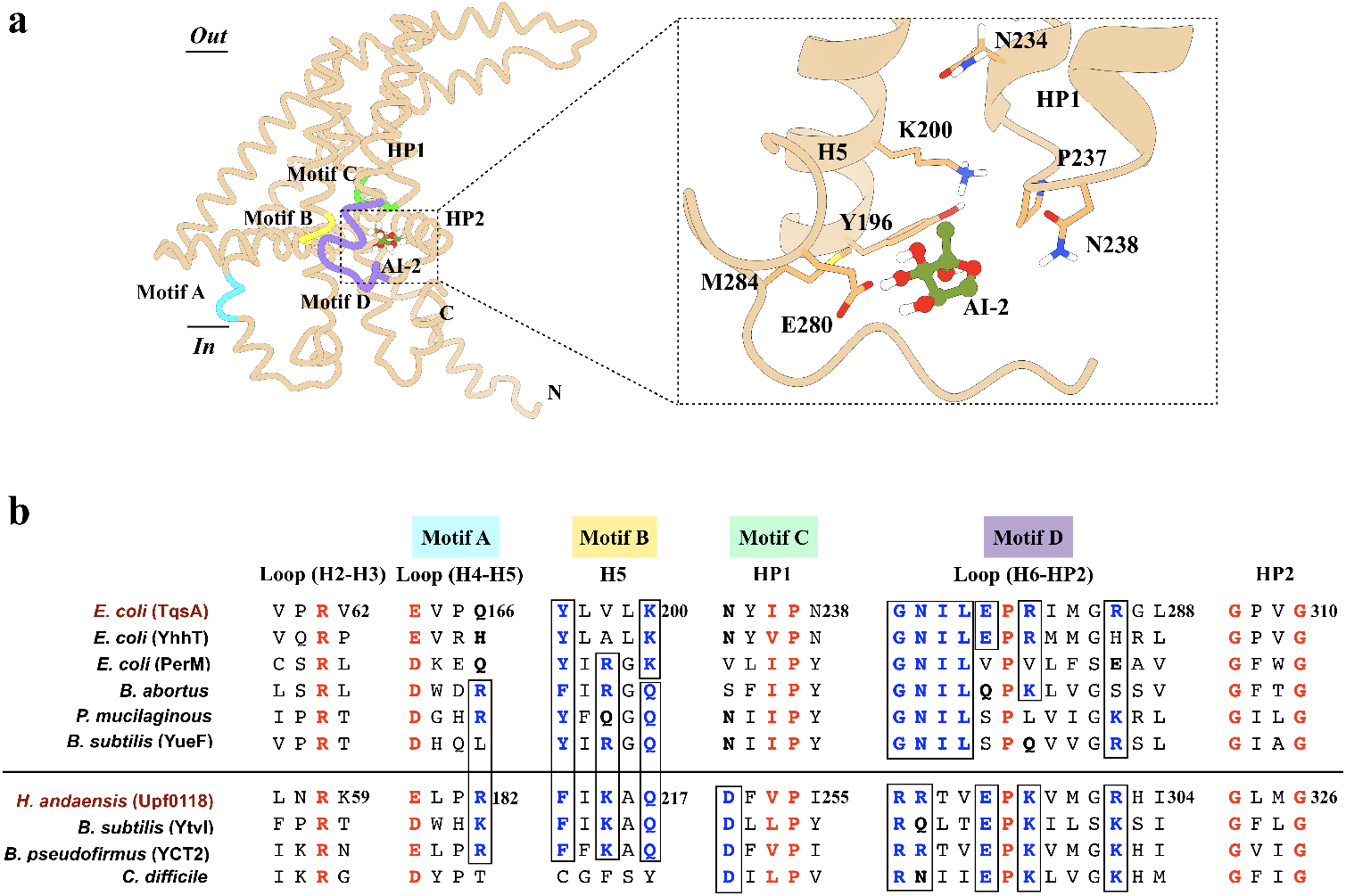
Molecular docking of AI-2 into the TqsA protomer. **a:** The AI-2 molecule is depicted in an olive-green color. A closer look on the residues lining the docked site is shown on the right. The predicted binding site is formed by the residues from H5, the tip of HP1 and the loop connecting H6 to HP2. Motifs which were previously recognized as critical for Upf0118 function were positioned on the TqsA monomer structure based on a sequence alignment. Motifs in the structure are shaded in distinct colors, Motif A: sky-blue, Motif B: yellow, Motif C: green, Motif D: purple. **b:** Multiple sequence alignment highlighting the relation between AI-2 exporters and Upf0118. Highly conserved residues are colored in red, and partially conserved ones in blue. The alignment is sectioned based on conserved residues and Upf0118 motifs. Homologs considered for the AI-2 exporter group include AI-2 exporters from *E. coli* (TqsA, YhhT and PerM), *Brucella abortus* (Uniprot: A0A7U8JIH7), *Paenibacillus mucilaginosus* (Uniprot: F8FLY5) and *Bacillus subtilis* (YueF, Uniprot: O32095). Similarly, homologs considered for Upf0118 group include proteins from *H. andaensis* (Upf0118, Uniprot: A0A1W5X0D5), *B. subtilis* (YtvI, Uniprot O50485), *Bacillus pseudofirmus* (YCT2, Uniprot: Q04454), *Clostridium difficile* (Uniprot: Q188A3). The choice of the homologs considered for multiple sequence alignment is based on the sub-groups within the AI-2 exporter family mentioned in the Upf0118 study^28^. YdiK from *E. coli* is not included in the alignment due to low sequence similarity. It constitutes a distinct subgroup in the family.

Recently, studies on Upf0118 from *H. andaensis* have shown that this Na^+^/H^+^ antiporter, together with its homologs, shares five functionally critical motifs (Motif A-E)^28^. Based on the sequence alignment, we explored the positioning of those motifs in the TqsA protomer structure (Fig. 5). Our results show that the potential substrate binding site in TqsA is surrounded by the motifs B, C and D, which are located on H5, the tip of HP1 and the loop connecting H6 to HP2, respectively. In the Upf0118 family proteins, motif B includes a lysine (Lys215^Upf0118^), followed one residue later by a polar glutamine residue (Gln217^Upf0118^). Mutagenesis studies have shown that Gln217^Upf0118^ is crucial for the Na^+^/H^+^ antiporter activity, while Lys215^Upf0118^ may be involved in the response of the antiporter activity to pH^28^. The motif B of TqsA also harbors a potentially positively charged lysine (Lys200^TqsA^), however, at the equivalent position of Gln217^Upf0118^ (Fig. 5 b). In TqsA, the motif C forms the short turn region of HP1 and is defined by a pattern of one small hydrophobic residue (Ile236^TqsA^) followed by one absolutely conserved proline (Pro237^TqsA^) as shown in Fig. 5 b. Asp251^Upf0118^ within this region is the most critical residue, and its substitution with even glutamic acid resulted in the complete loss of activity^28^. This aspartate residue is not present in the motif C of TqsA and other AI-2 exporters under study, but we could observe a polar residue, Asn234^TqsA^, at an equivalent position in TqsA and some of the other proteins (Fig. 5 b). The motif D in Upf0118 possesses two potentially positively charged arginine residues (Arg292^Upf0118^ and Arg292^Upf0118^), which were shown to be critical for the antiporter activity^28^. In TqsA, both arginines are replaced by a glycine and an asparagine residue, respectively (Fig. 5 b). Notably, a short E-P-R/K tripeptide can be found in the motif D of many AI-2 exporters as well as in the Upf0118 family proteins (Fig. 5 b). Glu296^Upf0118^ was shown to be vital for the function of Upf0118, and our docking studies also suggested the involvement of an equivalent residue in TqsA (Glu280^TqsA^) in substrate binding. In addition to the above-mentioned motifs, two other motifs (A and E) are identified in Upf0118. There are some highly conserved residues among the family including Glu163^TqsA^ present in motif A as shown in Fig. 5 b, which are located rather far from the putative substrate interaction site and their functional roles require further analyses. Motif E was not considered to be essential for the Na^+^/H^+^ antiporter activity of Upf0118 protein, and it is anyhow not present in other AI-2E family proteins as well. Taken together, the canonical AI-2 exporters, in particular TqsA from *E. coli*, share similar yet distinct motifs with Upf0118 family proteins. The potential substrate binding site of TqsA lacks several conserved charged or polar residues that are important for the Na^+^/H^+^ antiporter activity seen in Upf0118 family proteins, providing key evidence for the two proteins to belong to the same protein family but with different functions.

In addition to the docking experiment, native mass spectrometry (native-MS) was also employed to investigate the binding of AI-2 to TqsA. The mass spectrum of TqsA in 0.05% n-dodecyl-N, Ndimethylamine-N-oxide (LDAO) in the absence of AI-2 substrate display a charge state distribution corresponding to pentameric TqsA, which further confirms the oligomeric state that we observed in the structure (Fig. S11 a). Additionally, the spectrum also displayed a range of endogenous lipids copurified with the protein, whose masses are ~1388 Da, ~2963 Da and 4416 Da. Further lipidomic analysis on the TqsA sample showed the presence of common E. coli lipids PE (phosphatidylethanolamine), PG (phosphatidylglycerol) and CDL (cardiolipin) (Fig. S12 d). We then added 60 μM DPD/AI-2 to 4 μM TqsA and recorded the data. The resultant spectra display a clear shift (~140 kDa) in the peak positions for pentameric TqsA (Fig. S11 a), indicating the binding of this non-borate form of AI-2 to the TqsA complex. To further confirm the binding of AI-2 to TqsA, we analysed the samples in the delipidating detergent tetraethylene glycol monooctyl ether (C8E4), where the majority of the protein was found to be in monomeric form. When AI-2 was added, the mass spectra clearly show additional peaks which correspond to the binding of AI-2 to the TqsA monomer (Fig. S11 b). A similar methodology was also adopted for YdiK and we could observe AI-2 interaction with YdiK monomer as well (Fig. S11 c). Taken together, we could observe the interaction of AI-2 with TqsA both in silico and in vitro. However, further studies are required to elucidate the functional importance of the interacting residues.

### Discussion

Members of the AI-2 exporter superfamily are putative secondary active transporters that are widely distributed in prokaryotes and archaea^25^. This protein family (2.A.86) was renamed (formerly PerM family, 9.B.22) subsequent to the characterization of its first member TqsA from *E. coli* as an exporter of the quorum sensing signaling molecule autoinducer-2^31^. In this work, we could show that three AI-2 exporters from *E. coli* (TqsA, YdiK and YhhT) assemble into a homo-pentameric complex (Fig. 2 and Fig. S2). A pentameric assembly is a rare occurrence among membrane transporters. To the best of our knowledge, the only other transporter family with pentameric members is the formate/nitrite transporter (FNT) family^32^. In addition to the *E. coli* proteins, we also purified one another AI-2 exporter from the hyperthermophilic bactrerium *Aquifex aeolicus* (Uniprot id: Aq_740, O66948), which is the only family member (~ 40.2 kDa) present in its genome. Our preliminary data show that Aq_740 also forms a pentamer (Fig. S5). The 6.4 Å low resolution 3D reconstruction showed that it adopts a similar helical arrangement as the other characterized members (TqsA and YdiK) and it also substantiates that the pentameric state is a general characteristic of the AI-2 exporter superfamily (Fig. S5).

In this study, using cryo-electron microscopy, we have determined the first structures of two AI-2 exporters from *E.coli*, TqsA and YdiK at 3.35 Å and 2.80 Å overall resolution, respectively (Fig. 2 a, c). Both protein complexes comprise a central hydrophobic pore facing the cytoplasm, which extends around one-third into the membrane and holds an overall bowl-shaped appearance, opening up in the periplasmic side (Fig. 4 a, b). The presence of the pore connecting the periplasmic basin to the interior of the cell may insinuate that this region of the protein can potentially serve as a channel. However, the pore appears to be sealed by the hydrophobic filter in the case of TqsA and the presence of lipids in YdiK (Fig. 4 a).

Very recently, DeepMind has developed the artificial intelligence (AI) deep learning system AlphaFold for the prediction of protein structures^33^. During the preparation of this manuscript, the AlphaFold protein structure database was launched^34^, which also included the predicted structures of the four AI-2 exporters from *E. coli*. We compared our cryo-EM structure of TqsA with the one predicted by AlphaFold. The superimposition of both structures showed a surprisingly high similarity with an RMSD (root mean square deviation) of 3.8 Å (Fig. 6 a). The main differences are within the connecting loops and the region that is believed to undergo structural rearrangements necessary for transport. In the case of YdiK, our experimentally determined partial structure also shares a remarkable similarity with the corresponding portion of the AlphaFold structure as indicated by an RMSD of 4.6 Å (Fig. 6 b). Importantly, the predicted structure helped to fill in the missing parts of the experimental structure. It should be mentioned that the missing region of H3 and H4 was predicted with low confidence, which suggests a high flexibility and may explain why the density here cannot be well resolved. In addition, both YhhT and PerM were predicted to comprise around six helical segments and two helical hairpins and to share similar fold characteristics with TqsA (Fig. S10). Taken together, AlphaFold’s computational predictions appear to be quite noteworthy and accurate, at least for the four AI-2 exporters investigated here, although the prediction of the oligomeric state and topology currently remains not possible.

**Fig. 6.**
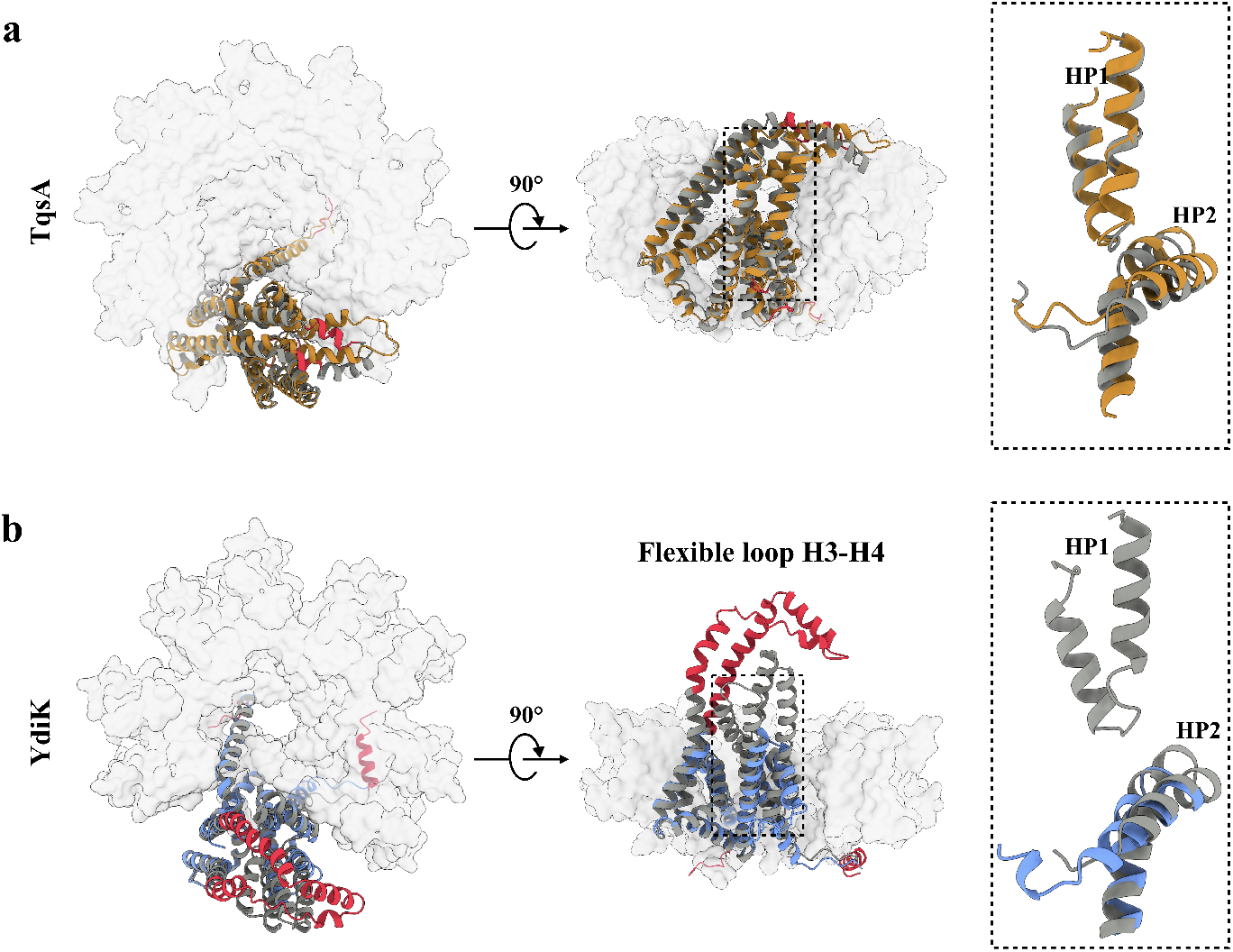
Comparison between cryo-EM and AlphaFold predicted structures of AI-2 exporters: **a, b:** Two orthogonal views of superimposition of the cryo-EM and AlphaFold predicted monomer structures of TqsA (**a**) and YdiK (**b**). The monomer structure is shown as a ribbon diagram in the pentameric complex. Cryo-EM structures of TqsA and YdiK are shown in brown and blue respectively. The predicted structures are shown in grey and low confidence areas with a per-residue confidence score lower than 70 are highlighted in red. The magnified view of the superimposed helical hairpins is shown on the right for both proteins.

Overall, the AI-2 exporter TqsA comprises several interesting structural elements. To begin with, the helix arrangement of H1 and H2 is absolutely peculiar (Fig. 4 a, b). It gives a perception of a “camera shutter” capable of widening and narrowing during the transporter activity. Another captivating feature in each protomer is the presence of two “helical hairpins” which is a characteristic and functionally critical structural element reported in many other transporters, such as the glutamate transporter SLC1, the nucleoside transporter CNT and the citrate transporter NaCT^35–38^ (Fig. S8 b). However, both hairpins in the AI-2 exporter are arranged distinctively relative to the other families (Fig. S8 b). In most of the other transporters, the tips of the helical hairpins face each other in the overall structure. In AI-2 exporter, HP1 is perpendicular to the membrane plane while HP2 is curved and bends almost parallel to the membrane plane facing the interior towards the periplasmic basin (Fig. S8 b).

All above-mentioned transporters possessing two helical hairpins employ the elevator-type mechanism for transport of their respective substrates^35,36,38–40^. Several common features among such transporter families have been highlighted in previous reports like the presence of two distinct domains, one rigid scaffold domain and one mobile transport domain normally segregated by a tilted helix supporting the major conformational changes during the transport cycle^39^. Our structures also reveal the presence of two such domains that are separated by a long and extremely bent helix H4 (Fig. 3 b, d and Fig. S8 a).

In addition, most elevator-type transporters exist as either homodimers or homotrimers, in which the subunit contacts are mainly mediated by the scaffold domain ^39^. Although AI-2 exporters assemble in an even higher oligomeric state, still the major contact points between the protomers are formed by the scaffold domains (Fig. S8 a). Moreover, a membrane topology with inverted repeats is another commonality of the elevator-type transporters. In TqsA, internal pseudosymmetry can be only observed in the C-terminal domain (Fig. 3 b).

The topology of the C-terminal transport domain of TqsA shows a remarkable similarity with that of sodium/potassium dependent glutamate and amino acid transporters, *e.g*., Glt_ph_ from *Pyrococcus horikoshii* and the human ASCT2^36,37^ (Fig. 7 a). Previous structural studies of glutamate transporters revealed that this protein forms a bowl-shaped trimer^36^. In both proteins Glt_ph_ and TqsA, the transport domain comprises a transmembrane helix followed by two helical hairpins and a connecting helix between the hairpins (Fig. 7 a). Although the aqueous basin in both proteins faces the same direction, the overall topology of the transport domain and the transmembrane helix connecting N- and C-terminal halves is inverted. This inversion event might be attributed to the fact that the scaffold domain of TqsA has one α-helix less compared to Glt_ph_, which also evolved to allow transport of the substrate in the opposite direction (Fig. 3 b). Glutamate transporters are responsible for the uptake of glutamate from extracellular fluids, while TqsA acts as an exporter of AI-2 to the periplasm.

**Fig. 7.**
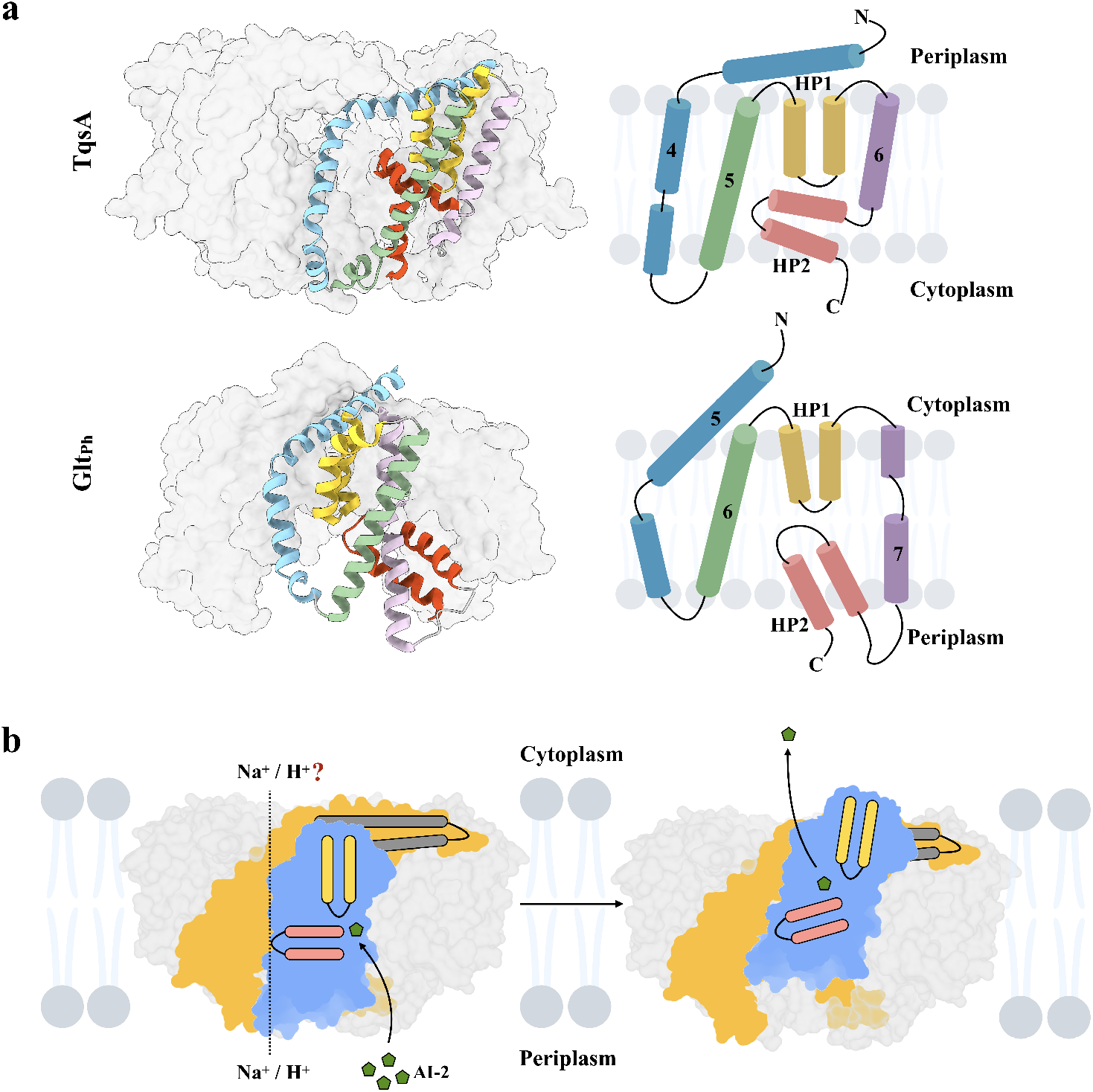
Proposed transport mechanism for AI-2 export. **a:** Similarities between the transport domain of AI-2 exporter TqsA (PDB: xxxx) and glutamate transporter (Glt_ph_) from *Pyrococcus horikoshii* (PDB: 2NWL) are shown. Individual segments of the transport domain of TqsA are shaded in different colors (H4 in blue, H5 in green, HP1 in yellow, H6 in purple and HP2 in red). Similarly, corresponding structural segments in Glt_ph_ are colored in a similar manner. The topology map of the transport domains of both proteins are included on the right of the figure for a better comprehension. **b:** Proposed transport mechanism of TqsA. AI-2 is depicted in an olive-green color in the figure and homo-pentameric AI-2 exporter TqsA is shown embedded in the membrane bilayer as viewed from the membrane plane. The lipids surrounding TqsA are depicted in faint blue. The scaffold and transport domains of the monomer are shown in yellow and blue, respectively. HP1 and HP2 are depicted in yellow and red, respectively and the long anti-parallel tilted helix bundle (H3 and H4) is represented in dark gray. The likely outwards facing conformation via elevator motion of TqsA is suggested in the figure on the right. The coupling ion for the transport is kept as an unresolved factor in the figure.

The observation of the structural similarity between the transport domain of the AI-2 exporter TqsA and that of glutamate transporter yields a strong clue for anticipating the transport mechanism of the AI-2 exporter. Glt_ph_ utilizes an elevator-type mechanism for providing alternating access of the binding site to either side of the membrane^36^. Each protomer of Glt_ph_ works as an independent transport unit. The binding site of glutamate in Glt_ph_ is observed at the interface of the two helical hairpins, also involving residues from the surrounding helices (TM7 and TM8) and loops in the transport domain. As shown earlier, a similar binding site was predicted for AI-2 in TqsA from our molecular docking studies involving residues from HP1, H5 and the loop connecting H6 to HP2 (Fig. 5 a). In Glt_ph_, HP1 is reported to be involved in substrate binding while HP2 acts as a gate, alternatively sealing and opening the binding site during the transport cycle. HP2 in the glutamate transporters also harbors a sodium ion filter, GXXG in prokaryotes and SXXS in eukaryotes^36,41^. The GXXG motif can be observed in HP2 of TqsA as well and these two glycine residues are absolutely conserved in the AI-2 exporter family (Fig. 5 b). The two highly tilted helices in the glutamate transport along which its transport domain slides, determine the distance the binding site can travel across the membrane during the transport cycle (fixed barrier)^39^. Similarly, in TqsA the two long tilted helices (H3 and H4) and the loop connecting H4 to H5 on the cytoplasmic side may act as a determinant or fixed barrier, deciding the distance which the transport domain can progress (Fig. S6 c). The flexible loop connecting H6 to HP2 also seems to be a participant in this barrier, but this speculation requires support from further studies (Fig. S6 c). Additionally, the large aqueous basin formed in the glutamate transporter is considered beneficial for allowing rapid access of the transmitter to the binding site during synapses^36^. A similar bowl-shaped architecture in TqsA might allow rapid diffusion of the pheromone AI-2 into the environment through the aqueous periplasmic bowl (Fig. S9).

Taken together, we suggest a similar transport mechanism for AI-2 exporters as exercised by glutamate transporters (Fig. 7 b). Each protomer can work as an independent transport unit adopting an elevator-type movement. In addition, we could observe a peculiar protein conformational change for Aq_740 in both 2D and 3D class averages during the single particle cryo-EM data processing (Fig. S5). One protomer in the pentameric complex was seen to be risen towards the periplasmic side. This observation supports our suggestion about the elevator-type mechanism for AI-2 exporters. However, further studies are required for a better understanding of the transport mechanism of this family.

In summary, many questions remain open, requiring further studies to elucidate the transport mechanism of the AI-2 exporters. These include the movement and gating mechanism of the hairpins, sodium being the coupling ion for the transport, the possibility of the presence of any ion channel and the identification of other potential substrates. Whereas proteins such as TqsA comprise a short α-helix at the C-termini, larger homologs in AI-2 exporter family contain a C-terminal ATP/GTP binding domain, a sensor GAF domains or a Tpr-like ligand binding domain^25^. In search of other potential substrates, such as cyclic AMP, cyclic GMP, nucleosides and even other quorum sensing signaling molecules including AI-3, an *in vitro* transport assay system has to be established^42^. So far, we were unable to elicit transport activity from TqsA using a number of methods, but the attempts are still ongoing. In this study we successfully characterized several members of the Autoinducer-2 exporter family, highlighted the structural uniqueness. Based on the structural analysis and comparisons, we could suggest a transport mechanism for this transporter family. This study on the pentameric AI-2 exporters will prove to provide benefits for future studies in the fields of quorum sensing, biofilm formation and antibiotic resistance.

## Materials and methods

### Cloning, expression and purification of autoinducer-2 exporters

Primers were designed based on the full-length gene sequences for each of the four selected AI-2 exporter family proteins from *Escherichia coli* K-12 (TqsA: [UniProt ID: P0AFS5], YdiK: [UniProt ID: P0AFS7], YhhT: [UniProt ID: P0AGM0] and PerM: [Uniprot ID: P0AFI9]). All primers are listed in the Supplementary Table. S1. Genomic DNA from the *E. coli* strain DH5α was isolated using the QuickExtract DNA extraction solution (Lucigen). The coding sequences of the four autoinducer-2 exporter genes were amplified using the Phusion DNA polymerase (Thermo Fisher Scientific). The resulting DNA fragments were gel-extracted and cloned individually into the pBADC3 plasmid using InFusion EcoDry cloning kit (TaKaRa). All genes were fused with a C-terminal StrepII-tag for affinity purification.

For protein overexpression, *E. coli* Top10 cells transformed with the expression vectors were grown in lysogeny broth (LB) medium supplemented with 100 μg/ml carbenicillin at 37°C to an optical density at 600 nm (OD600) of 0.5-0.6. Production of autoinducer-2 exporters was induced by the addition of 0.02% (w/v) L-arabinose, and incubation was continued for 4 h. Cells were harvested by centrifugation and stored at −80°C.

For protein purification, cells were resuspended in the lysis buffer (20 mM Tris-HCl, pH 7.5, 250 mM sucrose, 150 mM choline chloride) supplemented with protease inhibitors (1 mM phenylmethylsulfonyl fluoride, 1 mM benzamidine hydrochloride and 1 mM 6-aminocaproic acid), DNase I (approximately 50 μg/ml of buffer), 2.5 mM MgCl_2_ and 1 mM dithiothreitol (DTT). Cell disruption was conducted by passing through a microfluidizer at 8,000 psi and 4°C for six passes. The cell debris was removed by centrifugation at 4°C and 12,000 × *g* for 45 mins using a GSA rotor. Subsequently, the supernatant containing the membranes was centrifuged at 4°C and 43,000 × *g* for 2.5 hours. The pelleted membranes were resuspended in the lysis buffer without any additives to a concentration of around 20 mg total protein per ml of buffer. Aliquots of the membranes were flash-frozen in liquid nitrogen and stored at - 80°C. The total protein concentration was determined using Bradford protein assay.

All purification steps were performed at 4°C. The membrane proteins (100 mg total protein) were solubilized in the solubilization buffer (100 mM Tris-HCl, pH 7.5, 25% glycerol, 300 mM NaCl and 2% [w/v] n-dodecyl-ß-D maltoside [DDM]) with gentle stirring at 4°C for 1.5 hours. Following solubilization, the insoluble membrane fraction was removed by ultracentrifugation at 55,000 × *g* for 1 hour. The supernatant containing the solubilized proteins was supplemented with 40 μg/ml avidin and subsequently loaded onto a 5-ml StrepTrap HP column (GE Healthcare), which was pre-equilibrated with binding buffer (50 mM Tris-HCl, pH 8, 100 mM NaCl, 1 mM EDTA and 0.03% [w/v] DDM), with a peristaltic pump. Following loading, the column was washed with 20 column volumes (CV) of washing buffer (50 mM Tris-HCl, pH 8, 100 mM NaCl, 1 mM EDTA and 0.1% [w/v] glycol-diosgenin [GDN]). The StrepII-tagged proteins were eluted with 6 CV of the washing buffer supplemented with 5 mM desthiobiotin. The protein was further purified via gel filtration using Superose 6 Increase 10/300 GL column (GE Healthcare) pre-equilibrated with the purification buffer (50 mM Tris-HCl, pH 7.5, 150 mM NaCl, 0.006% [w/v] GDN). The proteins were concentrated with 50 kDa MWCO Amicon filters (Merck) to around 3-4 mg/ml for subsequent structural studies.

### SDS- and BN-PAGE

The purified proteins were analyzed by sodium dodecyl sulfate-polyacrylamide gel electrophoresis (SDS-PAGE) using 4-12% Bis-Tris NuPAGE gels (Invitrogen). Prior to SDS-PAGE analysis, protein samples were heated in the presence of 10 mM DTT at 90°C for 10 mins. For Western-blot analysis, proteins from SDS-PAGE gels were transferred onto a nitrocellulose membrane using iBlot blotting system (Thermo Scientific). Membranes were blocked for 1 h in TBST (10 mM Tris/HCl, 150 mM NaCl, 0.05% Tween 20, pH 8) containing 2% (w/v) biotin-free bovine serum albumin (BSA). The Strep-II-tagged proteins were immunodetected using the alkaline phosphatase-conjugated Strep-Tactin (IBA) and the color reaction was developed with 5-bromo-4-chloro-3-indolyl phosphate (BCIP) and nitroblue tetrazolium (NBT). The Blue native-PAGE was performed using the Novex 4-16% Bis-Tris gels (Invitrogen) according to the manufacturer’s instructions.

### Differential scanning fluorimetry

Differential scanning fluorimetry (DSF) measurements were performed with a Nanotemper Prometheus NT.48 (NanoTemper Technologies) at a heating rate of 1°C/min from 20°C to 95°C. Protein samples with a concentration ranging between 0.5 and 1.0 mg/ml were measured in triplicates in quartz capillaries (Prometheus NT.48 Series nanoDSF grade standard capillaries, NanoTemper Technologies). Fluorescence measurements were made with an excitation wavelength of 280 nm. The unfolding transition was monitored by detection of changes in the intrinsic tryptophan fluorescence ratio (F350/F330).

### Sample preparation and cryo-EM data collection

Before proceeding to cryo-EM data collection, protein samples (0.01 mg/ml) were negatively stained with uranyl formate and analyzed by negative staining electron microscopy using a Tecnai Spirit transmission electron microscope (TEM) operated at 120 kV. For cryo-EM, an aliquot of 4 μl of purified proteins (TqsA, YdiK and YhhT) at around 3-5 mg/ml concentration was applied to freshly glow-discharged C-flat holey carbon grids (R1.2/1.3, Cu, 300 mesh). The grids were blotted at blotting force 20 for 4 s at 4°C and 100% humidity and plunged into liquid ethane using a Vitrobot (Thermo Scientific).

High-resolution cryo-EM imaging was performed using a 300 kV Titan Krios G3i microscope (Thermo Scientific) equipped with a BioQuantum imaging filter (Gatan) and a K3 direct electron detector (Gatan). Cryo-EM data were collected in electron counting super-resolution mode at a nominal magnification of 105,000 ×. For TqsA, a total of 5,452 movies were collected with a total dose of 100 e^-^/ Å^2^ (100 frames) using the aberration-free image-shift (AFIS) correction in EPU. For YdiK, a total of 7,378 movies were collected with the same exposure settings. In addition, 989 movies were collected for YhhT with a total dose of 43 e^-^/ Å^2^ (40 frames). The defocus range used for data collection was from −1.1 to −2.1 μm, and cryoSPARC live was employed for constantly monitoring the quality of the incoming movies and for the on-the-fly data processing^43^.

### Single particle data processing

For TqsA, dose-fractionated and gain-corrected movies were subjected to beam-induced motion correction using MotionCorr2^44^. Afterwards, motion corrected micrographs were contrast transfer function (CTF) estimated using CTFFIND4^45^. 3,314 micrographs with CtfMaxResolution values better than 4 A° were selected for further processing. Particle picking was done with CrYOLO1.5 using a general model for low pass filtering^46^. Around 1.2 million particles were picked from selected micrographs and imported in Relion3.1^47^. The particles were extracted with a down-sampled pixel size of 3.348 Å and a box size of 80 pixels. These particles were subjected to multiple rounds of reference-free 2D classification, which resulted in 427,253 particles. Particles from several best 2D classes were used for the initial model building and a 3D classification was applied. 133,640 particles from the best 3D classes were extracted with the original pixel size of 0.837 Å per pixel and a box size of 320 pixels. For the 3D refinement, C5 symmetry was imposed because the pentameric organization was clearly visible in the 2D class averages as well as in 3D maps as shown in the Supplementary Figs. S2 b and S3. Refinement was also performed separately with C1 symmetry as well. After the first refinement step, C5 and C1 symmetry EM maps were generated at 3.72 Å and 4.25 Å resolution, respectively (Fig. S3). Iterative rounds of particle polishing and CTF refinement were performed to improve the quality of both maps. Finally, C5 and C1 symmetry EM maps of global resolutions of 3.35 Å and 3.99 Å were obtained, respectively, according to the gold standard FSC = 0.143 criterion. The polishing step was carried out with the first 70 fractions, filtering out the later dose-damaged fractions. Local resolution maps were generated using Relion3.1, as shown in the Supplementary Fig. S2 c, d. The data processing workflow is illustrated in the Supplementary Fig. S3.

For the YdiK dataset, a similar strategy was applied, and a detailed flowchart is shown in Supplementary Fig. S4, and local resolution maps are shown in Fig. S2 e, f. Finally, a C5 symmetry map at 2.80 Å global resolution was obtained from 619,311 particles.

### Model building and validation

For both TqsA and YdiK, initial partial models were obtained using a combination of ARP/wARP 8.0 and Buccaneer in the CCP-EM software suite ^48,49^. These models were corrected and extended *de novo* into the EM density maps using COOT^50^. A combination of phenix.real_space_refine^51^ and Refmac^52^ was used for the restrained refinement for improving the fitting of the models into the EM maps^52,53^. Map to model and cross validation was carried out using Phenix suite and MolProbity^54,55^. Refinement and validation statistics for both models are summarized in Table 1.

**Table. 1.** Cryo-EM data collection, refinement and validation statistics.

| Specifications | TqsA (EMDB: xxxxx)<br>(PDB:xxxx) | YdiK (EMDB: xxxxx)<br>(PDB:xxxx) |
| --- | --- | --- |
| **Data collection and processing** |  |  |
| Magnification | 105,000 | 105,000 |
| Voltage (kV) | 300 | 300 |
| Electron exposure (e<sup>-</sup>/Å<sup>2</sup>) | 80 | 80 |
| Defocus range (μm) | -1.1 to -2.1 | -1.1 to -2.1 |
| Pixel size (Å) | 0.837 | 0.831 |
| Symmetry imposed | C5 | C5 |
| Initial particle images (no.) | 1,218,552 | 1,222,739 |
| Final particle images (no.) | 134,300 | 619,311 |
| Map resolution (Å) | 3.35 | 2.80 |
| FSC threshold | 0.143 | 0.143 |
| Map resolution range (Å) | 3.0 - 7.2 | 2.6 - 6.5 |
| **Refinement** |  |  |
| Initial model used | - | - |
| Model resolution (Å) | 2.9 | 2.5 |
| FSC threshold | 0.143 | 0.143 |
| Map sharpening B factor (Å<sup>2</sup>) | -30 | -10 |
| **Model composition** |  |  |
| Non-hydrogen atoms | 13020 | 8410 |
| Protein/nucleotide residues | 1700 | 1105 |
| Ligands | - | - |
| **B factors (Å<sup>2</sup>) (min/max/mean)** |  |  |
| Protein | 18.22/197.18/109.91 | 71.51/175.24/100.68 |
| Ligands | - | - |
| **R.m.s deviations** |  |  |
| Bond lengths (Å) | 0.010 | 0.006 |
| Bond angles (°) | 1.282 | 1.002 |
| **Validation** |  |  |
| MolProbity score | 2.13 | 1.90 |
| Clashscore | 13.42 | 9.94 |
| Poor rotamers (%) | 0.49 | 0 |
| CaBLAM outliers (%) | 2.38 | 2.39 |
| **Ramachandran Plot** |  |  |
| Favored (%) | 91.72 | 94.42 |
| Allowed (%) | 8.28 | 5.58 |
| Disallowed (%) | 0 | 0 |

### Native mass spectrometry

Prior to MS analysis, the sample buffer was exchanged to 200 mM ammonium acetate, pH 8.0, and 2 times the critical micelle concentration (CMC) of various detergents (namely LDAO and C8E4) using a Biospin-6 column (BioRad) and introduced directly into the mass spectrometer using gold-coated capillary needles (prepared in-house). Data were collected on a Q-Exactive UHMR mass spectrometer (Thermo Scientific). The instrument parameters were as follows: capillary voltage 1.1 kV, quadrupole selection from 1,000 to 20,000 m/z range, S-lens RF 100%, collisional activation in the HCD cell 200 V, trapping gas pressure setting at 7.5, temperature 200°C, In-source trapping 300 V, resolution of the instrument 6250. The noise level was set at 3 rather than the default value of 4.64. Data were analysed using Xcalibur 4.2 (Thermo Scientific). AI-2/ DPD was provided by Dr. Rita Ventura’s group, ITQB Portugal^56^.

### Substrate docking

Docking of AI-2 to the TqsA structure was performed with Glide^57^. The initial coordinates of AI-2 were taken in the cyclic form from the LsrB structure (PDB ID 6DSP) and an ensemble of conformations was generated using Ligprep (Schrödinger Release 2020-1: LigPrep, Schrödinger, LLC, New York, NY, 2020). Docking was carried out over a search space of the entire protein volume with 75 runs. In each run a box size of 15 × 15 × 15 Å for the inner box and 45 × 45 × 45 Å for the outer box was used. The binding poses obtained from all docking runs were pooled and then clustered using the cluster module of Gromacs program (v2020.2) with the Jarvis-Patrick method and a cut-off of 3 Å^58^.

### Sodium ion/proton antiporter assays

Two methods for investigating the functional role of AI-2 exporters as Na^+^/H^+^ antiporters were conducted as described previously^27,59,60^. One of them, the growth complementation assay, was performed using the *E. coli* Knabc strain ^59^, which was generously provided by Prof. Etana Padan, (Hebrew University, Israel). Briefly, the cells were transformed individually with the expression vectors containing the genes encoding AI-2 exporters. Overexpression of the proteins was induced by the addition of 0.02% (w/v) L-arabinose for 3 hours and the growth of the cells was monitored in both liquid and solid media containing 200 mM NaCl. The expression levels of the proteins were determined by Western blot analysis. All cultures were carried out in duplicates and the study was done multiple times at different pH values to confirm the results.

Another assay inspects the Na^+^/H^+^ antiporter activity in everted membrane vesicles using the pH dependent fluorescent dye acridine orange^61^. Briefly, membrane vesicles of TqsA, NhaB and empty pBADC3 plasmid from Knabc transformants were prepared in a similar way as the membranes were prepared for purification of the proteins except that the cells were disrupted at lower pressures ~3000 psi. Acridine orange was used as a fluorescent probe for detection of sodium ion induced pH variations. Fluorescence was measured using a Hitachi F4500 fluorimeter at excitation and emission wavelengths of 495 nm and 530 nm, respectively. Everted vesicles (approximately 100 μg of total protein) were added to the buffer containing 10 mM MES-Tris pH 8, 145 mM choline chloride, 5 mM MgCl_2_ and 2 μM acridine orange. The vesicles were acidified by addition of 2.5 mM Tris-D-lactate and after stabilization of the fluorescence, dequenching was induced with 50 mM NaCl.

## Data availability

The atomic coordinates for TqsA and YdiK have been deposited to PDB under the accession numbers: XXXX (TqsA) and XXXX (YdiK). The EM density maps including the C1 maps and masks used for final processing for both TqsA and YdiK have been deposited to EMDB under the accession numbers: EMD-XXXXX (TqsA) and EMD-XXXXX (YdiK). Further information can be acquired from the corresponding author upon request.

## Acknowledgements

We would like to thank the staff at the EM facility at the Max Planck Institute of Biophysics, Simone Prinz, Dr. Susann Kaltwasser and Mark Linder for their support, training and assistance for microscopy data collection; Hannelore Mueller for technical assistance in the lab; Jiangfeng Zhao and Philipp Valina Allo for help in cloning and purification of Aq_740; Dr. Schara Safarian for initial training and assistance at the microscope; Dr. Yongchan Lee for guidance and useful insights for cryo-EM data processing; Dr. Rita Ventura’s group, ITQB, Portugal for providing synthetic AI-2; Prof. Victor Sourjik and Dr. Karina Xavier from MPI for Terrestrial Microbiology and Instituto Gulbenkian de Ciência respectively for all their help and swift replies. This work was supported by the Max Planck Society.

## Author contributions

RK designed the experiments, purified proteins, acquired and processed cryo-EM data, built atomic models and analyzed data. ARM performed molecular docking studies. JRB carried out native-MS measurements. JK performed mass spectrometry analyses. SW provided technical assistance for high resolution cryo-EM data collection. UE assisted for building atomic models. CM provided technical assistance in the laboratory. CV and GH provided the work setup. HX supervised the research and provided guidance for data interpretation; RK and HX prepared figures and wrote the manuscript. All authors discussed the results, read and approved the manuscript. H.M supervised and funded this work.

