## supplementary materials for "Cryo-EM structures of pentameric autoinducer-2 exporter from *E. coli* reveal its transport mechanism"

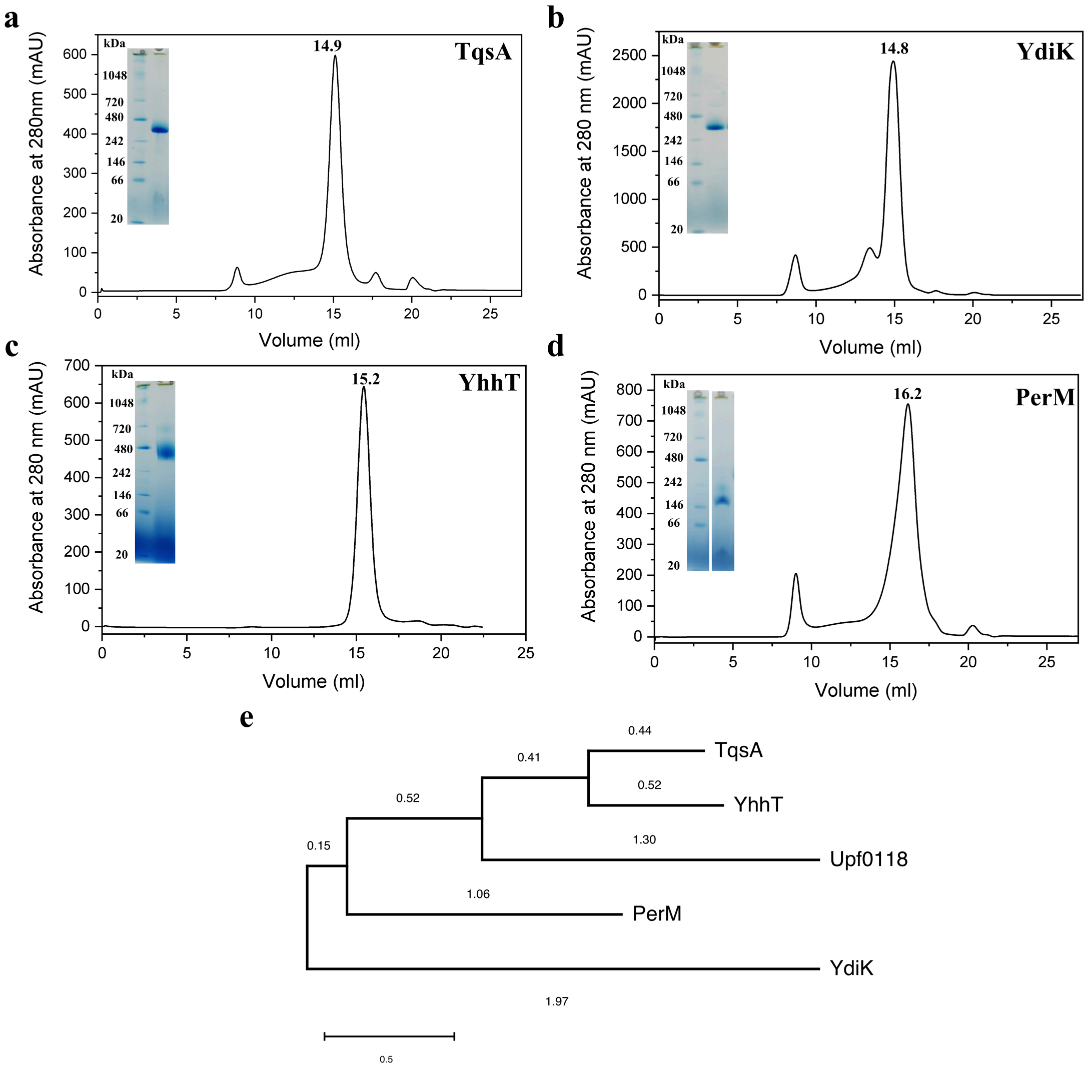


**Fig. S1 | Purification and phylogenetic analyses of AI-2 exporters from *E. coli*.** Size exclusion chromatography (SEC) and BN-PAGE profiles for the four AI-2 exporters from *E. coli*, TqsA **(a)**, YdiK **(b)**, YhhT **(c)** and PerM **(d)**. The retention volumes for each of the proteins are included. Superose 6 increase 10/300 GE column was used for the final size exclusion step. **e:** Phylogenetic analysis of the four AI-2 exporters and Upf0118 family using maximum likelihood method. MEGAX was utilized for the analysis and the branch distances are mentioned on the respective branches of the tree.

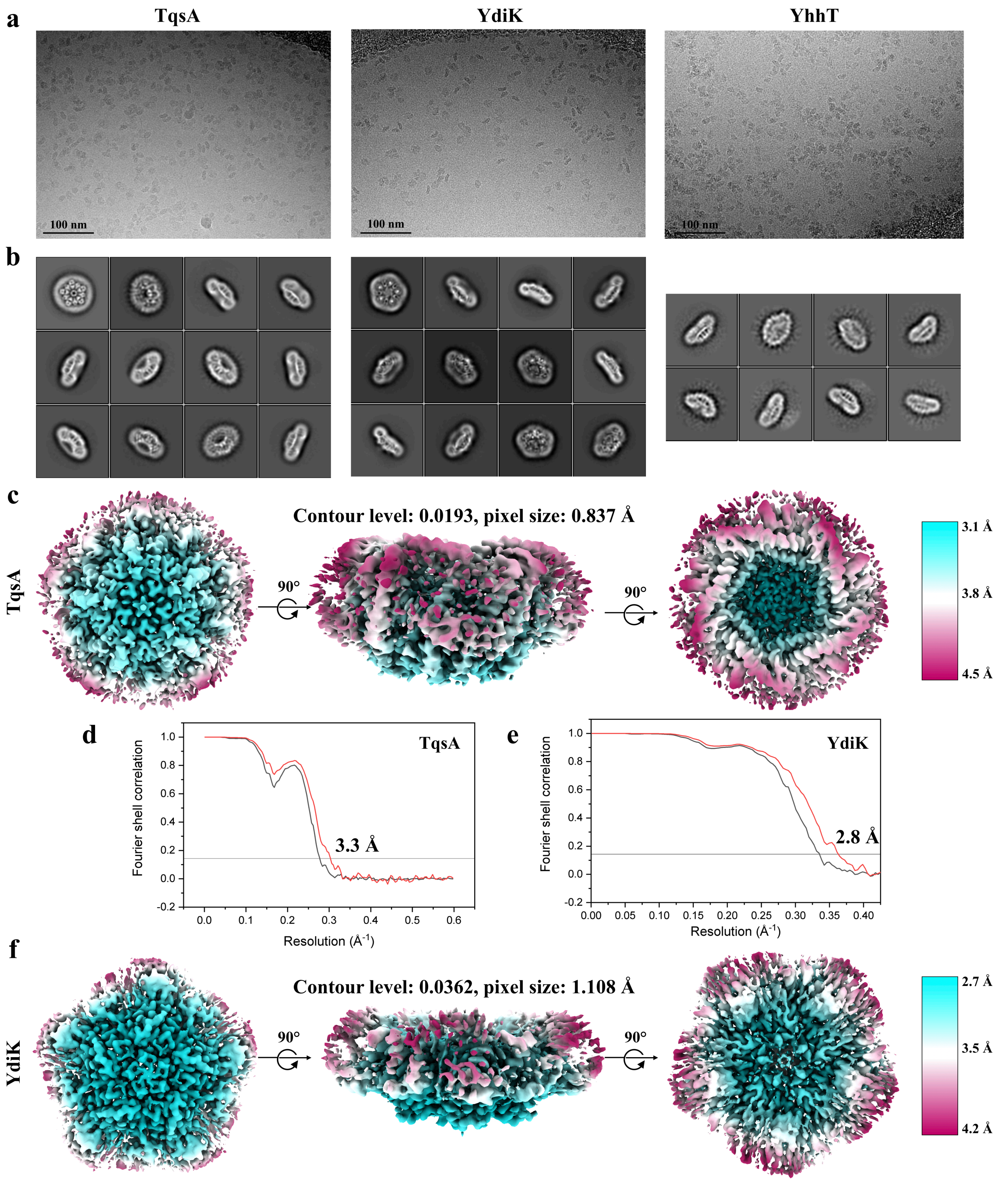


**Fig. S2 | Cryo-EM on AI-2 exporters. a:** Raw micrographs for the three GDN-purified AI-2 exporters TqsA, YdiK and YhhT from Titan Krios TEM at 300 kV. **b:** Reference-free 2D class averages for the proteins. **c, f:** Local resolution EM maps for TqsA and YdiK. **d, e:** Fourier shell correlation (FSC) curves for the AI-2 exporters TqsA and YdiK. The profiles for masked and unmasked maps are shown in red and black, respectively.

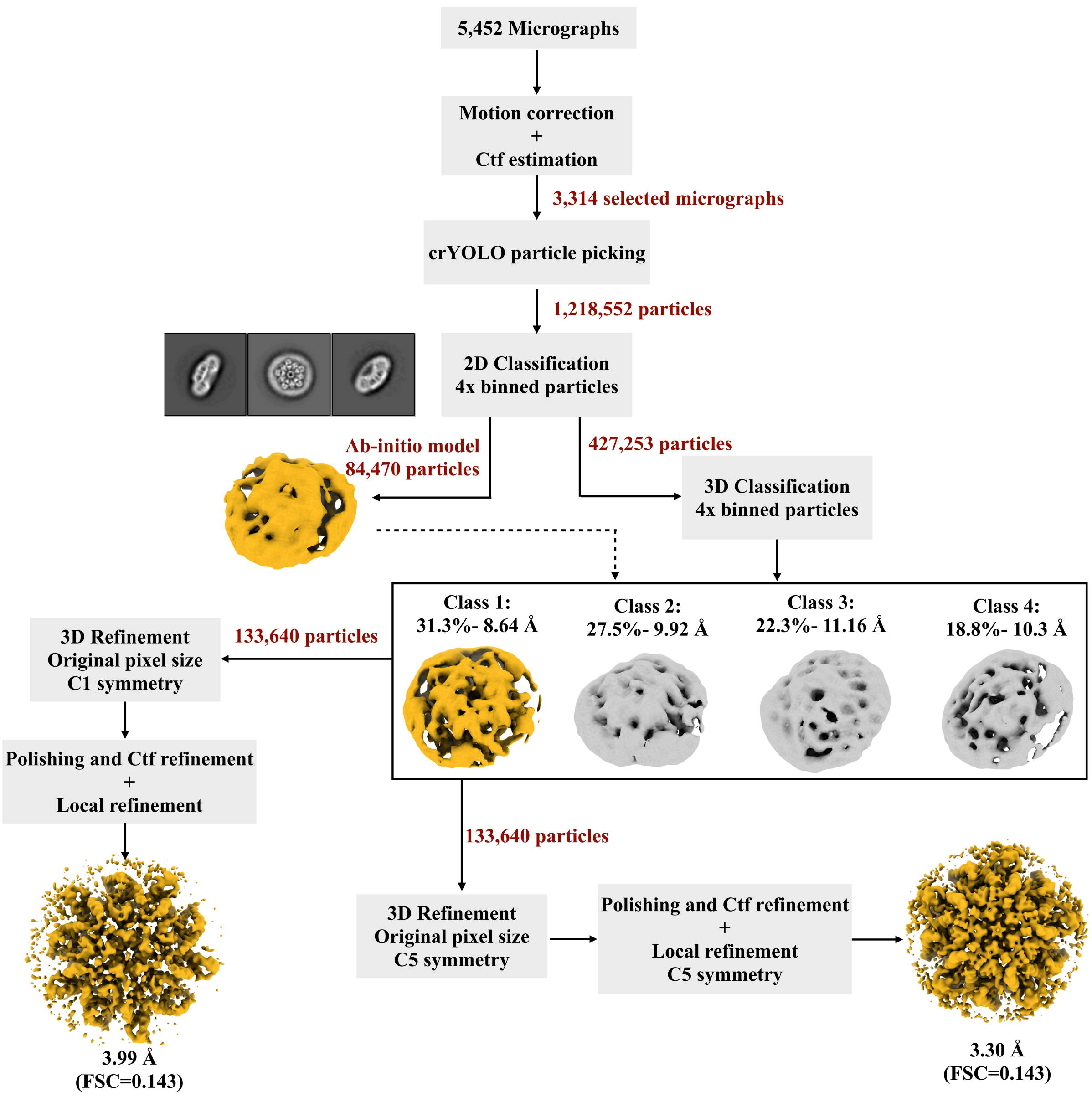


**Fig. S3 | TqsA single particle processing workflow.** Detailed single particle cryo-EM workflow adopted for TqsA dataset is described.

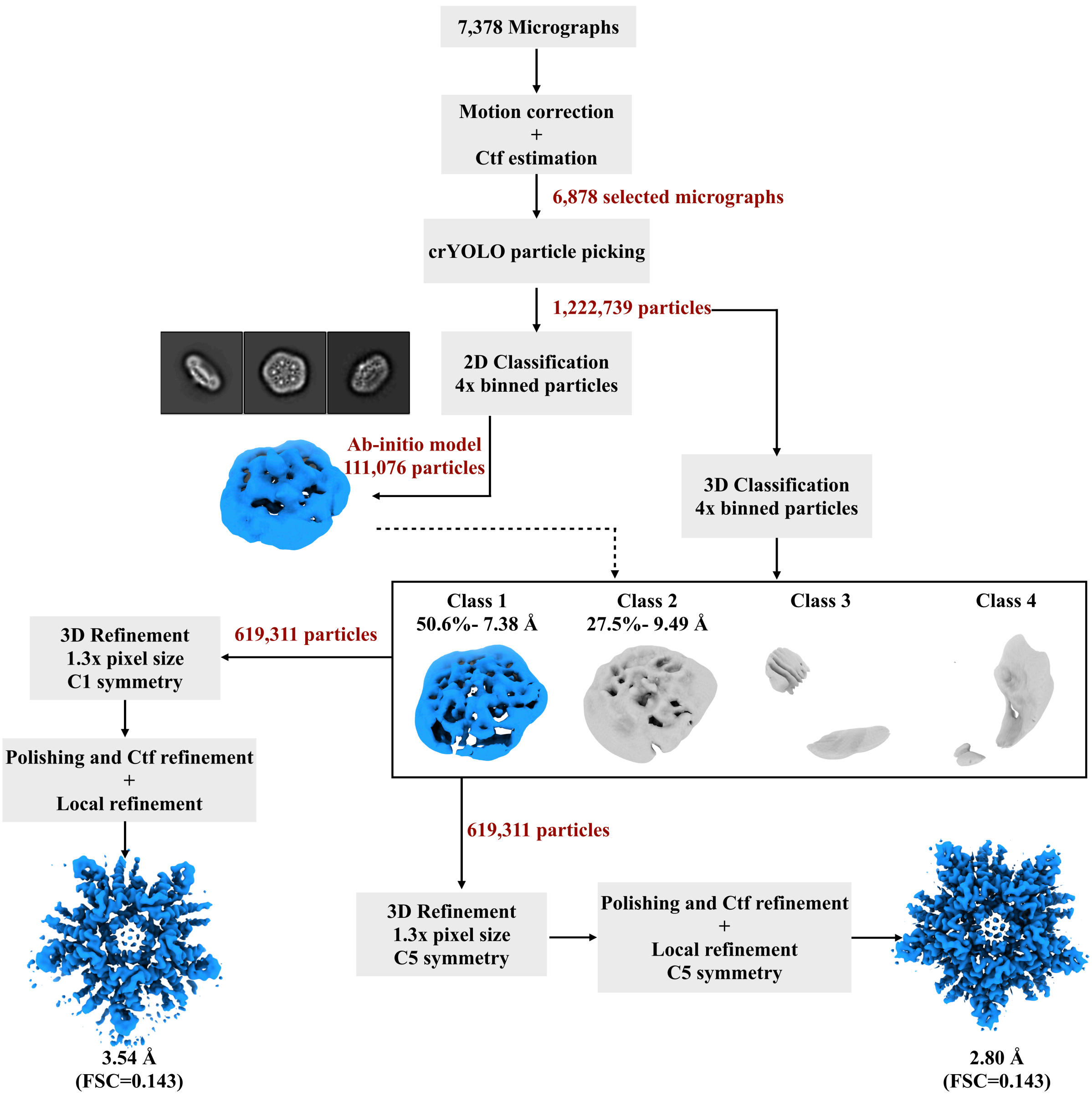


**Fig. S4 | YdiK single particle processing workflow.** Detailed single particle cryo-EM workflow adopted for YdiK dataset is shown.

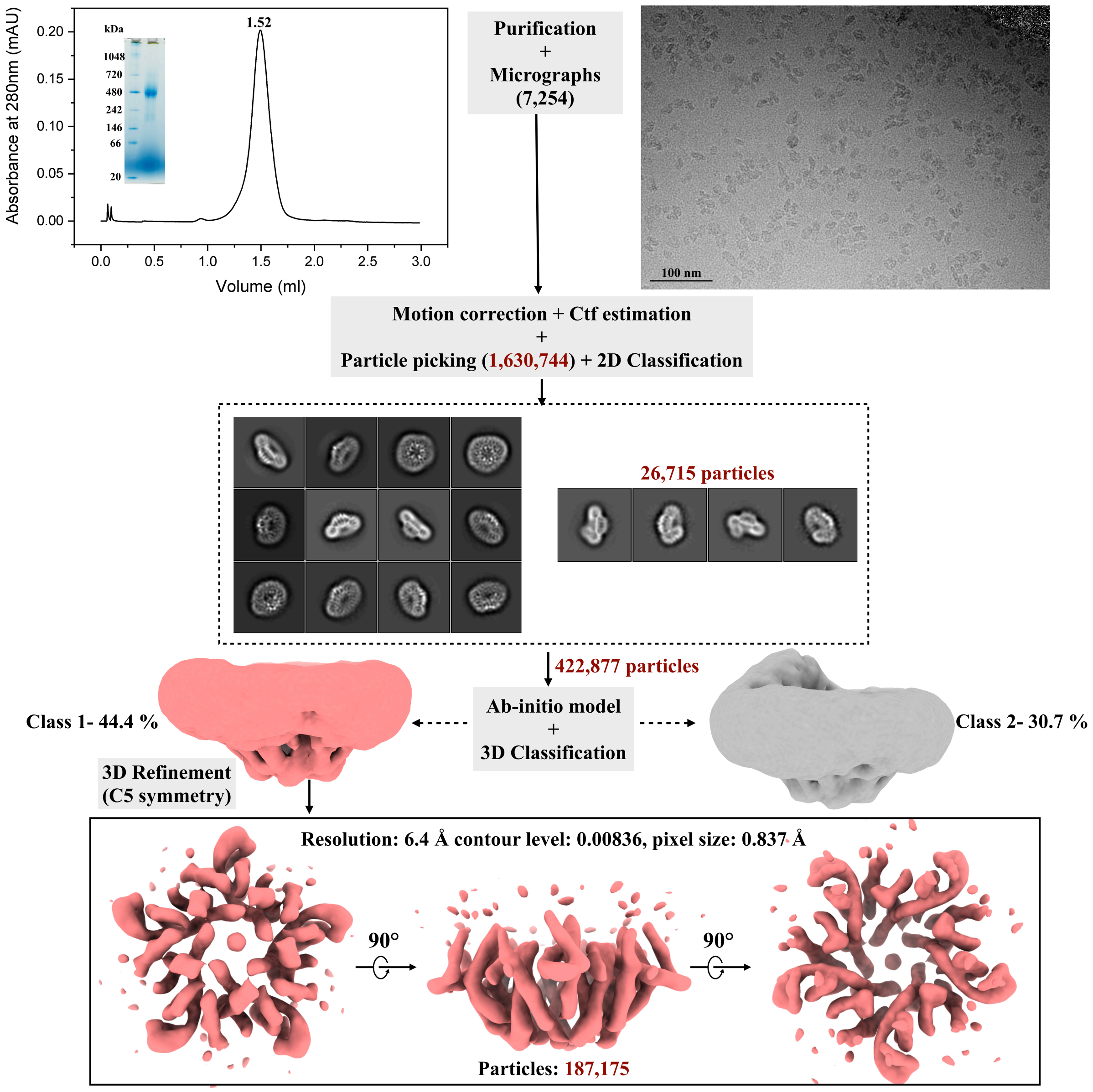


**Fig. S5 | Cryo-EM studies on Aq_740 from *A. aeolicus.*** A flowchart for the entire workflow adopted for Aq_740 is represented in the figure. Analytical SEC and BN-PAGE profile for Aq_740 are shown in the beginning on the top of the figure. Superose 6 3.2/300 GE column analytical column was used for the final size exclusion step. An illustrative raw micrograph for GDN purified AI-2 exporter Aq_740 from Titan Krios TEM at 300 kV with 0.837 Å pixel size is shown on the right. A total of 7,254 gain corrected movies were collected. Reference-free 2D class averages for Aq_740 were obtained from RELION 3.1 with an initial particle stack of around 1,630,744 particles. Later two different and almost equally populated 3D classes were obtained as shown. One of them depicted in light coral shade was further refined with imposed C5 symmetry. It yielded a low-resolution 3D reconstruction of 6.40 Å which is shown in different angular views at the bottom of the figure upon anti-clockwise rotation along the X-axis. The final particle stack of the map is around 187,175 particles.

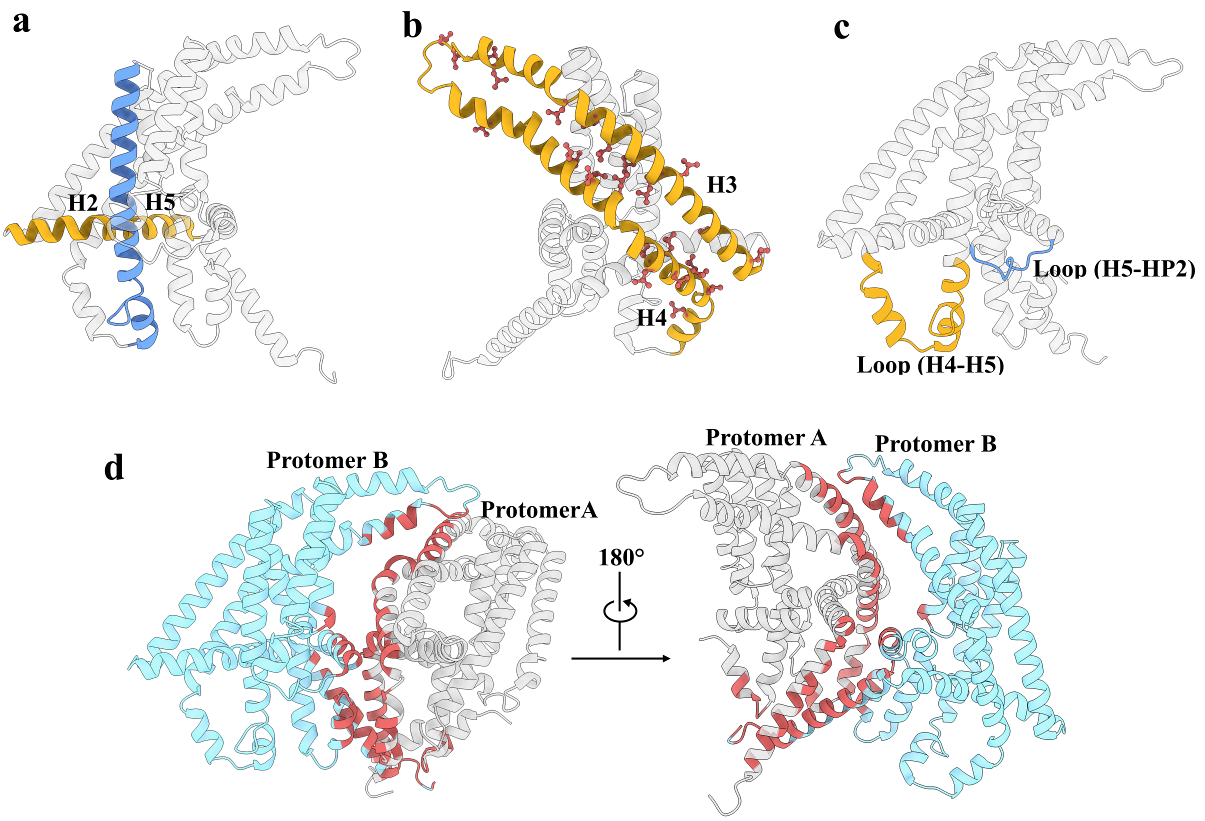


**Fig. S6 | Structural features of AI-2 exporter, TqsA. a:** Cross shaped architecture formed by helices H2 and H5 at the interface of scaffold and transport domains in TqsA is highlighted. Yellow is used for the scaffold domain (N-half) and blue for the transport domain (C-half). **b:** "Leucine zipper like motif" covering the tilted antiparallel helix bundle (H3 and H4) of the TqsA monomer is illustrated in this subsection. **c:** Loops suspected essential for the elevator motion of the transport domain in TqsA are focused on in the figure. **d:** The dimer interface within the homo-pentameric complex is highlighted in the figure for TqsA. The interacting residues are shown in red. The major interacting residues are located at the N- and C- terminus of TqsA. Loops connecting helices H3-H4 and H4-H5 also appear essential. The dimer interface and the interacting residues were calculated using PDBePISA. The estimated solvation free energy ΔG for the formation of the dimer interface is ~39.7 kcal/mol.

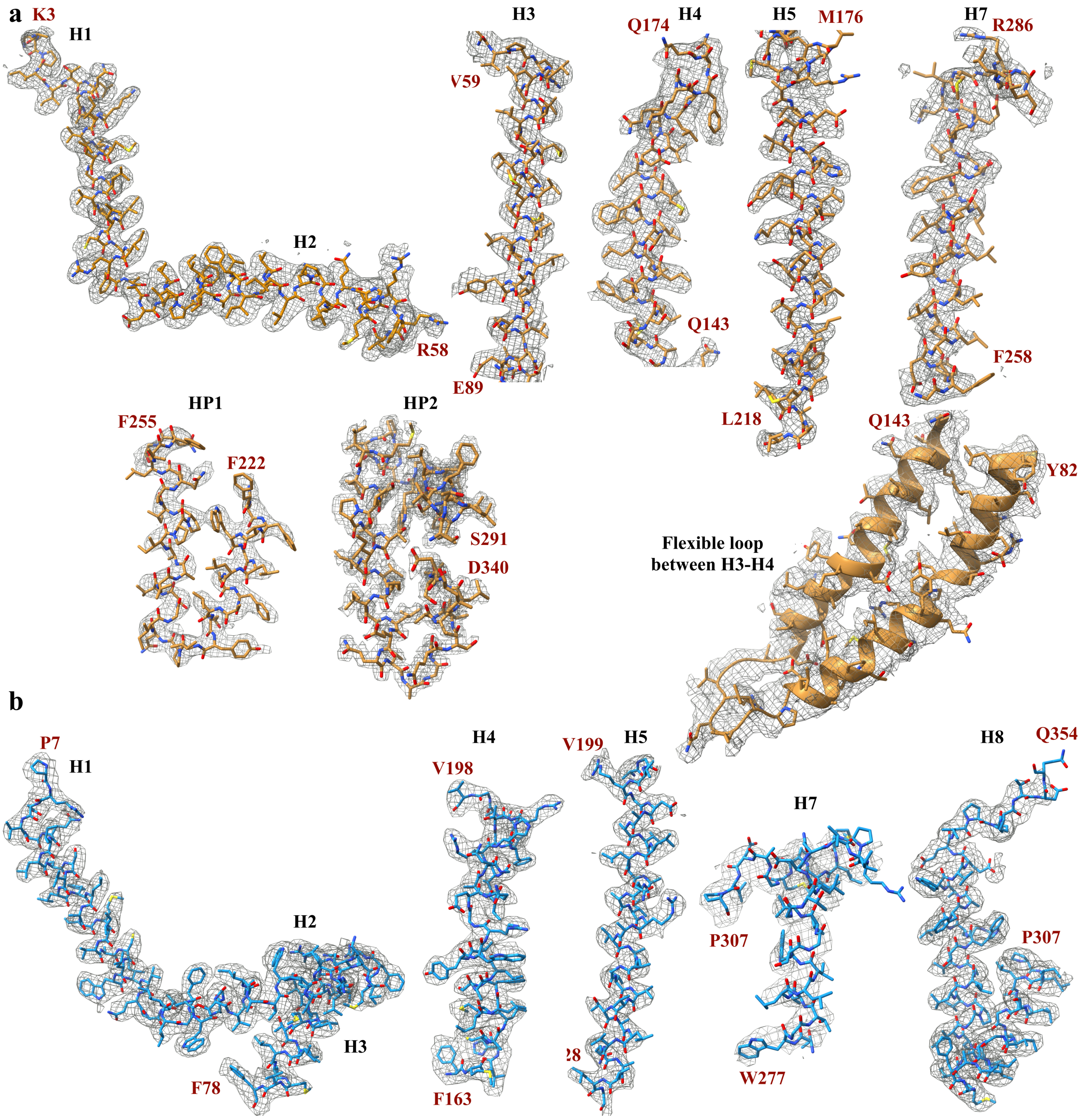


**Fig. S7 | Model fitting of the AI-2 exporters. a:** Model fitting of TqsA in the EM density map for different regions of the monomer structure. The residues constituting a particular structural region are indicated in red. **b:** Model fitting of YdiK in the EM density map for different regions of the monomer structure are shown with the similar nomenclature as that used for TqsA in a.

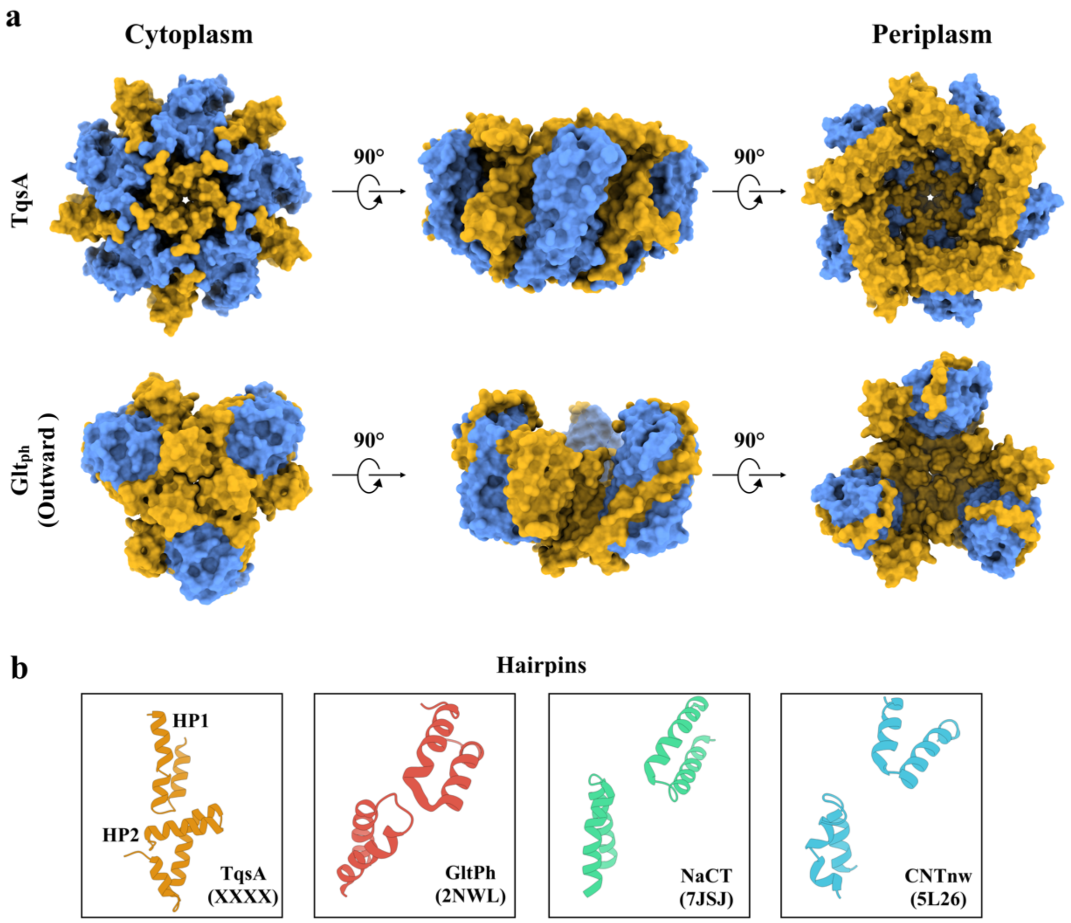


**Fig. S8 | Comparison between AI-2 exporter and other transporter families adopting elevator mechanism. a:** Contrast between AI-2 exporter TqsA (PDB: 7NB6) and glutamate transporter $Glt_{ph}$ (PDB: 2NWL) is shown. The scaffold domain in both the surface representative models are shown in yellow and the transport domain is illustrated in blue. **b:** The figure focuses on the difference of the helical hairpins in four different transporter families (AI-2 exporter (TqsA); XXXX, glutamate transporter; 2NWL, sodium citrate transporter; 7JSJ and nucleoside transporter; 5L26).

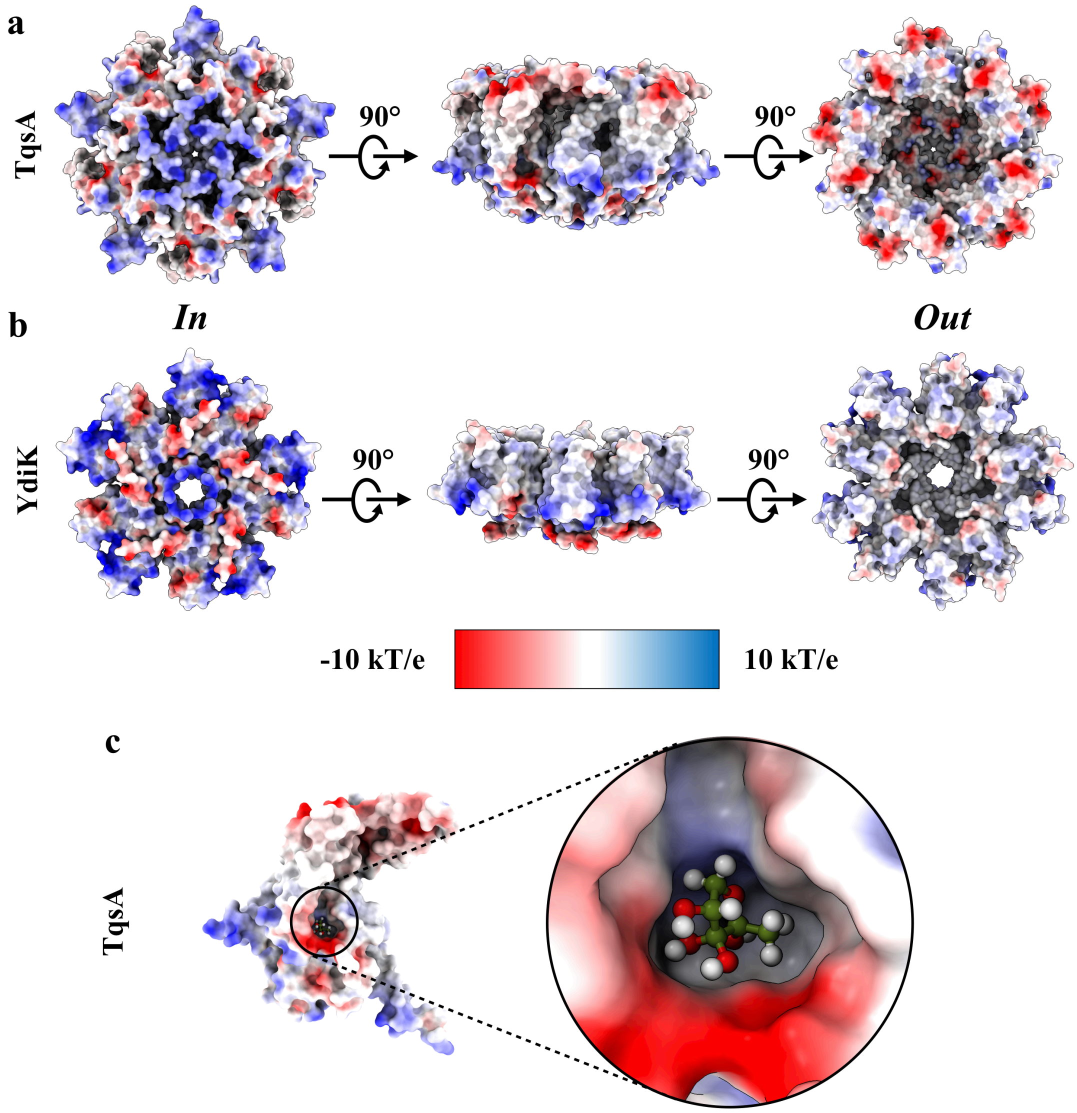


**Fig. S9 | Electrostatic potential surface maps of TqsA and YdiK.** Electrostatic surface potentials of TqsA (**a**) and YdiK (**b**) calculated by APBS are colored ranging from blue (+10 *kT/e*) to red (-10 *kT/e*). **c:** Close-up view on the charges around the docking site of AI-2 in TqsA.

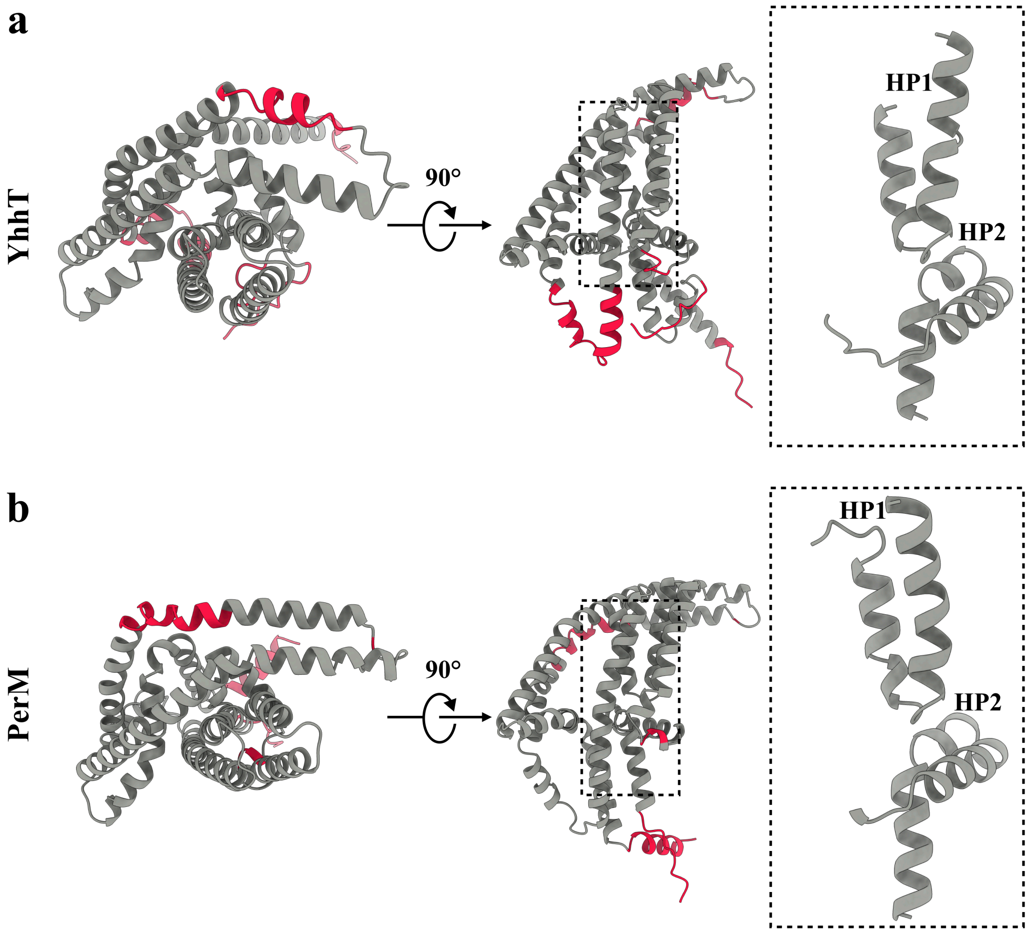


**Fig. S10 | AlphaFold predicted monomer structures of YhhT and PerM.** Both the predicted structures are shown in grey with two different views, **a:** YhhT and **b:** PerM. The low confidence areas in the structures are highlighted in red. The helical hairpins for both the predicted models are shown magnified on the right.

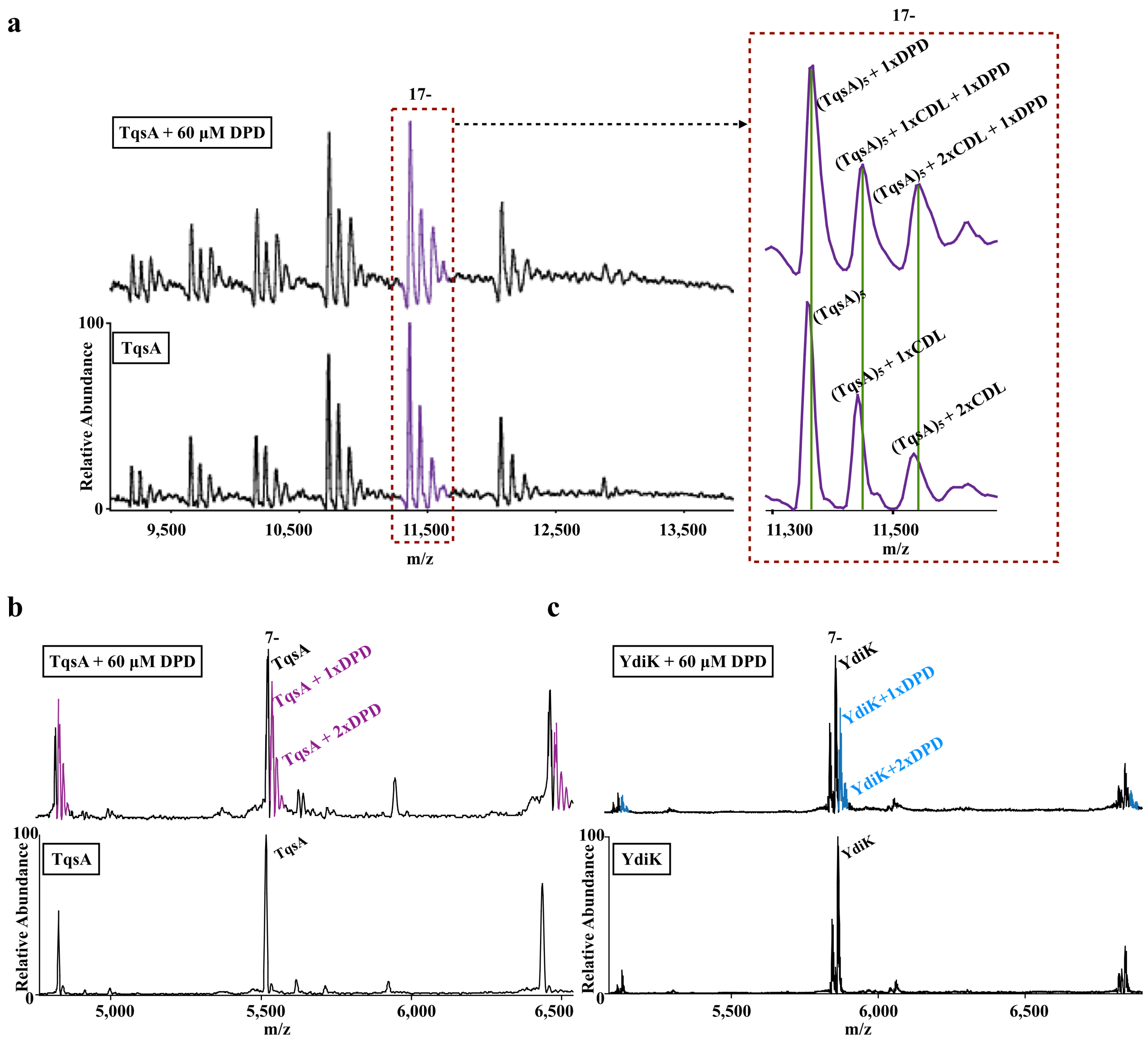


**Fig. S11 | Interaction analysis of AI-2 exporters with AI-2/DPD using Native-MS. a:** Two spectra for TqsA are shown in the figure, the upper one after the addition of DPD and the lower one devoid of any substrate. The peaks corresponding to the pentameric TqsA in both spectra are highlighted in purple. For clarity, a zoomed-in view of the 17- charge state is shown on the right as an inset. CDL in the figure represents a cardiolipin molecule. The binding of DPD to pentameric TqsA is evident from the shift in the peak centre **b:** The spectra of TqsA in C8E4 detergent with and without the addition of AI-2 show that the protein is majorly in the monomeric form and the additional peaks developed around it in the upper spectrum are highlighted in purple which indicates AI-2 interaction with the protein. **c:** The spectra of YdiK in C8E4 also showed a similar behavior that was observed in the case of TqsA in b, further confirming that YdiK also binds to AI-2 (highlighted in blue).

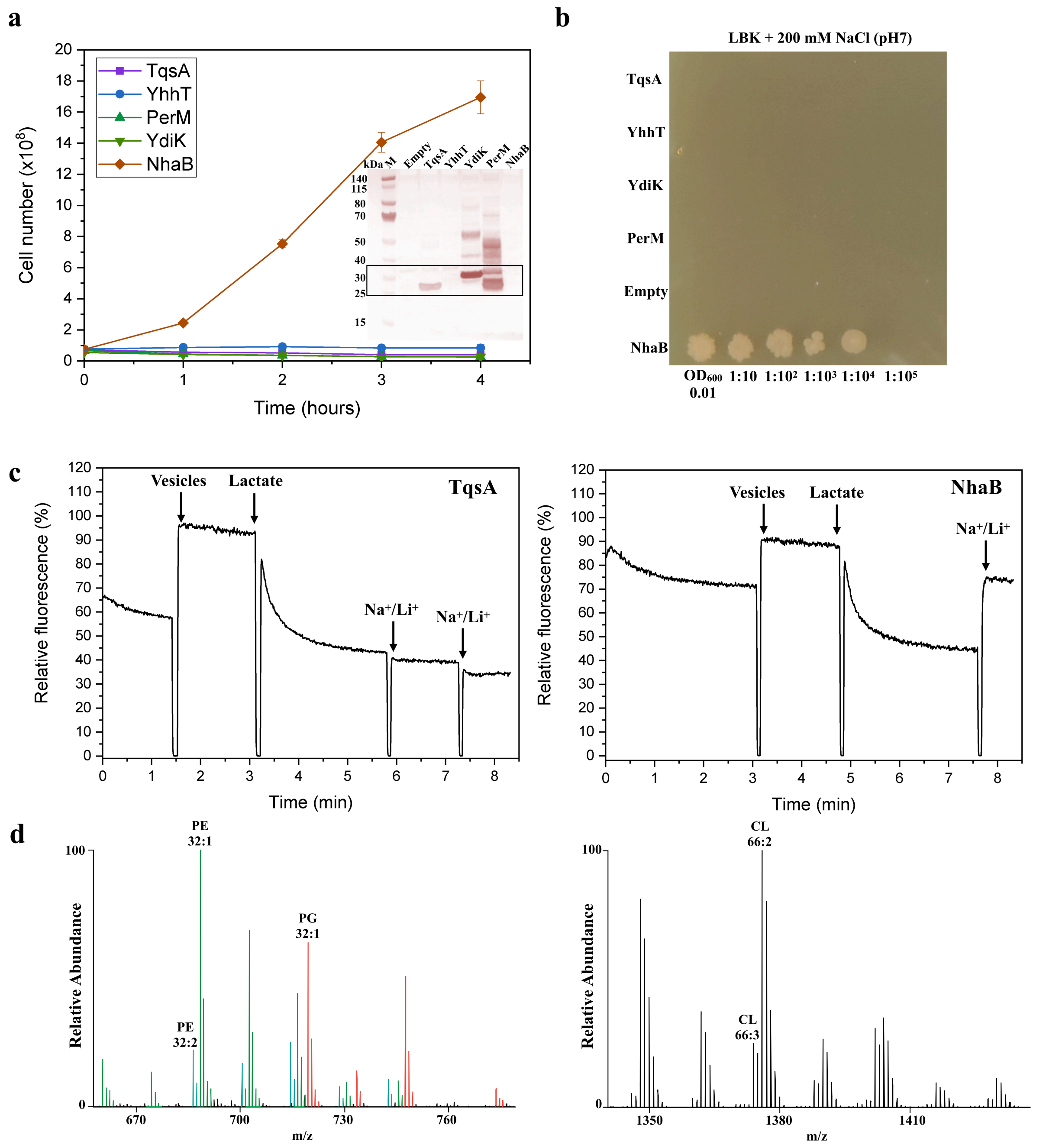


**Fig. S12 | Sodium proton antiporter assays and lipidomic analysis of purified TqsA. a:** Growth complementation sodium proton antiporter assay in LBK medium with 200 mM NaCl. The starting $OD_{600}$ of all the cultures was maintained around 0.06-0.07. Western blot analyses for the expression of the exporters are also included in the plot. *E. coli* NhaB couldn't be detected via Western blot either due to low expression levels or inaccessible Strep-II tag. **b:** Similar growth complementation sodium proton antiporter assay on LBK agarose medium with 200 mM of NaCl with *E. coli* NhaB as a positive control. **c:** Acridine orange assay profiles for AI-2 exporter TqsA on the left and the positive control

*Ec*NhaB on the right. **d:** Lipidomic analysis of GDN-purified TqsA. *E. coli* lipids, phosphatidylethanolamine (PE), Phosphatidylglycerol (PG) and cardiolipin (CL) were observed.

```
                                                 10        20        30        40
TqsA_Ec/1-344          ------------------------------MAKPIITLNGLKIVIM  16
YhhT_Ec/1-349          ------------------------------METPQPDKTGMHILLK  16
PerM_Ec/1-353          ----------------------------MLEMLMQWYRRRFSDPEA  18
BARG_01524_Ba/1-658    ----MVKKPTPARQTVTKTIRQSETIGRNGDIHVNVEVESFSGASV  42
YueF_Bs/1-369          ------------------------------MLKSKVHFWTLQILFV  16
F8FLY5_Pm/1-357        ---------------------------MDRFSKNSLLLWMVYMLLA  19
Upf0118_Ha/1-352       ---------------------------------MFRYLSKRQWILL  13
YtvI_Bs/1-371          ------------------------------MNQSYITIFFRTLFVI  16
Yct2_Bp/1-385          MNKVNKINAYTCSSFQLHTIMKISFYVVERGFELAAFFTRRTIWII  46
Q188A3_Cd/1-363        ----------------------MNEQDHTSYQNKQRFLINFCYLVV  24

                            50        60        70        80        90
TqsA_Ec/1-344          LGMLVIILCG-------IRFAAEIIVPFILALFIAVILNPLVQHMV  55
YhhT_Ec/1-349          LASLVVILAG-------IHAAADIIVQLLLALFFAIVLNPLVTWFI  55
PerM_Ec/1-353          IALLVILVAG----FGIIFFFSGLLAPLLVAIVLAYLLEWPTVRLQ  60
BARG_01524_Ba/1-658    RRQAGFWLGAMAFFVLFLVVFSSVLLPFVAGMALAYFLDPVADRLE  88
YueF_Bs/1-369          LLIIFVATKVSFVFQPFIVFISTLFFPMLIAGILYFIFNPVVRLL-  61
F8FLY5_Pm/1-357        LIILFMLQQLRPLFVGIYQFLREILTPFIIAMIISYVLNPVVTLLG  65
Upf0118_Ha/1-352       LLGILFVVAG--------YFILPVSVPLIIALITALFLNPAVRWMQ  51
YtvI_Bs/1-371          SMTAGSIAAA--------YYSFPLTYPFLIALILSSVIHPVVDYLD  54
Yct2_Bp/1-385          ISILLFLVAA--------YFILPVSLPLVAALLTALILTPAVNALQ  84
Q188A3_Cd/1-363        ILGLLMLLGK---------YAFSVLLPFVFAFLIASLLNKPVLYIS  61

                                100       110       120       130
TqsA_Ec/1-344          -RWRVPRVLAVSILMTIIVMAMVLLLAYLGSALNELTRTLPQYRNS 100
YhhT_Ec/1-349          -RRGVQRPVAITIVVVVMLIALTALVGVLAASFNEFISMLPKFNKE 100
PerM_Ec/1-353          -SIGCSRRWATSIVLVVFVGILLLMAFVVLPIAWQQGIYLIRDMPG 105
BARG_01524_Ba/1-658    -KVGLSRLAASIVILLIFLMILIIGLMIVVPILATQLADFISNLPG 133
YueF_Bs/1-369          -EKKIPRTLSILLIYLLFIGLLAFISASVGPIITAQVTGLFNNLPD 106
F8FLY5_Pm/1-357        -ARKVPRTIAVLLIYAVFITSTTVVLMNVIPMFVRQLAELNEHMPD 110
Upf0118_Ha/1-352       FRFRLNRKMAVTIVFLLFVIMIGLLGTYAVTRAVTQLVELADNAPS  97
YtvI_Bs/1-371          KVTGFPRTINVLGVLAFFLLAAFGVLTILVAEIVTGTAYLAKTLPP 100
Yct2_Bp/1-385          RKTKIKRNVAVMLVFTVFVVFIGLSGYYIATKAITQGTQIVENSPQ 130
Q188A3_Cd/1-363        EKLLIKRGIVATISVFLFFLIAGIIVSVIGTYLIYGIKEIFHFLPT 107

                         140       150       160       170       180
TqsA_Ec/1-344          IMTPLQALEPLLQ---------RVGIDVSVDQLAHYIDPNAAMTLL 137
YhhT_Ec/1-349          LTRKLFKLQEMLP---------FLNLHMSPERMLQRMDSEKVVTFT 137
PerM_Ec/1-353          MLNKLSDFAATLPR-------RYPALMDAGIIDAMAENMRSRMLTM 144
BARG_01524_Ba/1-658    YITQLQSLLANRDSEWLK---KYIGIDSTVIQQNLSSLLQQGAGFL 176
YueF_Bs/1-369          YIKQIQALTKDLS---------HSQWFTWMMNQDY-VSISKIEQSL 142
F8FLY5_Pm/1-357        VAYKAQSVVQNMN---------ELQFLPDSVRTGINQSLSKLENGI 147
Upf0118_Ha/1-352       YINQINNVLINWQN-NMN---SFTQNMPSEFVDKVSVELQNTIDTT 139
YtvI_Bs/1-371          HISTFISYCEKLFTTHIQPLYNELTLLFQELETNQQASIVTHIQTL 146
Yct2_Bp/1-385          YISDINRAWLNFQR-NLE---EKYENLPPELVQEINITVTNTLSDL 172
Q188A3_Cd/1-363        MFEELVIPLTETVI-------KEVELFSGSFNFSFIELIEANMPVL 146

                             190       200       210       220
TqsA_Ec/1-344          TNLLTQ----LSNAMSS---------------IFLLLLTVLFMLLE 164
YhhT_Ec/1-349          TALMTG----LSGAMAS---------------VLLLVMTVVFMLFE 164
PerM_Ec/1-353          GDSVVKIS--LASLVGLLTIAVY---------LVLVPLMVFFLLKD 179
BARG_01524_Ba/1-658    STLLQS----LWNSGKSLIDIAG--------LFVVTPVVAFYMLLD 210
YueF_Bs/1-369          TSFLQN----LPQNITSSLSAVFGVVTNITLVIITVPFILFYMLKD 184
F8FLY5_Pm/1-357        SLAIAN----YINSIGSTINTLF--------LIFIIPFVAFYILKD 181
Upf0118_Ha/1-352       TQTLSQKL--QLSNIAAFAAKIP----EYLISFLVYLIALFLFMLE 179
YtvI_Bs/1-371          GDSAAKNAGLLLSHILEMIPRFFALLPNTAAVLIFSLLATFFMTKD 192
Yct2_Bp/1-385          RSNISDRN--LIQDITSLISSIP----GYLVTFLVYLIALFLFMLE 212
Q188A3_Cd/1-363        MNEMSVFIVSLSNNIVSGITDIVSSVPLLFMKTIITIVASIFIAID 192
```

```
                                     240                 250                 260                 270
TqsA_Ec/1-344        VPQLPGKFQQMMAR--PVEGMAAIQRAIDSVSHYLVLKTAISIITG 208
YhhT_Ec/1-349        VRHVPYKMRFALNN--PQIHIAGLHRALKGVSHYLALKTLLSLWTG 208
PerM_Ec/1-353        KEQMLNAVRRVLPRN-RGLAGQVWKEMNQQITNYIRGKVLEMIVVG 224
BARG_01524_Ba/1-658  WDRMVNSIDSWVPRKQLHTVRRIAREMNAAVAGFIRGQGTLCLILG 256
YueF_Bs/1-369        GHRFPHLAVKILPASYRTEGLKIFKDLSDTLAAYFQGQLLICLFVG 230
F8FLY5_Pm/1-357      FQLLEKTTLAIVPRNHRKEIVSMLMDIDTALGNYIRGQFTVCMIVG 227
Upf0118_Ha/1-352     LPRLKDKMHGNFTESTSEKVKFMNARLSYVVFGFLKAQFLVSIVIF 225
YtvI_Bs/1-371        WHKLKAMLVLILPDRVTANSKAISSELKKAMTGFIKAQAVLVFITM 238
Yct2_Bp/1-385        LPRLREKLYSYLSERTKEKVNFMTSRLSYVIWGFFKAQFLVSIIIF 258
Q188A3_Cd/1-363      YPTIKRFIVLQIPKNKSYLLAEAKAFTIDTIIKCGFSYLLIFAITF 238

                       280                 290                 300                 310                 320
TqsA_Ec/1-344        LVAWAMLAALDVRFAFVWGLLAFALNYIPNIGSVLAAIPPIAQVLV 254
YhhT_Ec/1-349        VIVWLGLELMGVQFALMWAVLAFLLNYVPNIGAVISAVPPMIQVLL 254
PerM_Ec/1-353        IATWLGFLLFGLNYSLLLAVLVGFSVLIPYIGAFVVTIPVVGVALF 270
BARG_01524_Ba/1-658  TYYAIGLTLTGLNFGLLIGFFAGLISFIPYIGSFVGLALAIGVALV 302
YueF_Bs/1-369        TACFIGYLIAGLPYALILGIVMAITNIIPYVGPFLGAAPAVIVGFM 276
F8FLY5_Pm/1-357      LLAYIGYWLIGMPYALLLACVVAVFNIIPYLGPFLGAAPALIVAAT 273
Upf0118_Ha/1-352     VVCLIGLFWITPEVAIVMSLIIWIVDFVPIIGSIVILGPWALYMLI 271
YtvI_Bs/1-371        VIVFIGLSLLKVEHAATIAFLIGLVDLLPYLGAGSVFVPWILYLSI 284
Yct2_Bp/1-385        IVTLIGLLFIAPEVALLMAFIIWLIDFVPIIGSIVILAPWAIFQLI 304
Q188A3_Cd/1-363      SELYVGFVIINISYAGIIALFIALLDILPVLGTGSILIPWCIISLF 284

                                   330                 340                 350                 360
TqsA_Ec/1-344        FNG--FYEALLVLAGYLLINLVFGNILEPRIMGRGLGLSTLVVFLS 298
YhhT_Ec/1-349        FNG--VYECILVGALFLVVHMVIGNILEPRMMGHRLGMSTMVVFLS 298
PerM_Ec/1-353        QFG-AGTEFWSCFAVYLIIQALDGNLLVPVLFSEAVNLHPLVIILS 315
BARG_01524_Ba/1-658  QFWPDWIMVCTVAAVFFLGQFIEGNILQPKLVGSSVGLHPVWLMFA 348
YueF_Bs/1-369        DS---PAKALFAIIVVVIVQQLDGNLLSPLVIGKRLNTHPLTIILL 319
F8FLY5_Pm/1-357      IS---VKMVLFVILVNTACQIMEGNVISPQVVGRSLKMHPLFIIFA 316
Upf0118_Ha/1-352     VGD--IAMGGQLAMLAIILLAIR-RTVEPKVMGRHIGLSPLATLIA 314
YtvI_Bs/1-371        TGQ---LPQAIGIGILYLVVLIQRQLTEPKILSKSIGIDPLATLIA 327
Yct2_Bp/1-385        VGD--VSTGSKLLILAAVLLIIR-RTVEPKVMGKHIGLSPLATLIA 347
Q188A3_Cd/1-363      LGD---YSMALGIALLYIVITIIRNIIEPKLVGKHMELHPVLALAS 327

                        370                 380                 390                 400                 410
TqsA_Ec/1-344        LIFWGWLLGPVGMLLSVPLTIIVKIALEQTAGGQSIAVLLSDLNKE 344
YhhT_Ec/1-349        LLIWGWLLGPVGMLLSVPLTSVCKIWMETTKGGSKLAILLGPGRPK 344
PerM_Ec/1-353        VVIFGGLWGFWGVFFAIPLATLIKAVIHAWPDGQIAQE-------- 353
BARG_01524_Ba/1-658  LFAFGSLFGFTGMLVAVPAAAAVGVLVRFALNSYLRSPMYDPANNR 394
YueF_Bs/1-369        LIGAGSFGGILGMILAVPVYAVVKAFFLNIVRLIKLRQRSRLEENA 365
F8FLY5_Pm/1-357      LLVGGEIAGIAGLILAVPFFAVMKVILQHVFSYYIHRRTPT----- 357
Upf0118_Ha/1-352     MYIGLQLIGLMGFILGPLLVIAFNSAKEAGIIRWNFKL-------- 352
YtvI_Bs/1-371        LFAGFKLFGFLGLIAGPAVLVIIQAFITTGALKEIWSYITVQQK-- 371
Yct2_Bp/1-385        MYLGLMLFGVIGFIIGPLLVIAFTSAKEAGIIKLNFKL-------- 385
Q188A3_Cd/1-363      MLTGLHFFGFIGLFGIPLLIAFLKKLNDKEIIHILH---------- 363

                                420                 430                 440                 450
TqsA_Ec/1-344        ----------------------------------------------
YhhT_Ec/1-349        SRLPG----------------------------------------- 349
PerM_Ec/1-353        ----------------------------------------------
BARG_01524_Ba/1-658  SNPDAGPLIEAGDNTGREKWDGAKIRWDLMGAAMHDAPRQIPLNLE 440
YueF_Bs/1-369        KPAE------------------------------------------ 369
F8FLY5_Pm/1-357      ----------------------------------------------
Upf0118_Ha/1-352     ----------------------------------------------
YtvI_Bs/1-371        ----------------------------------------------
Yct2_Bp/1-385        ----------------------------------------------
Q188A3_Cd/1-363      ----------------------------------------------
```

```
                             470       480       490       500
TqsA_Ec/1-344        ----------------------------------------------
YhhT_Ec/1-349        ----------------------------------------------
PerM_Ec/1-353        ----------------------------------------------
BARG_01524_Ba/1-658  HQPGYNREDLIVTASNRAAVDLIDRWPNWLSPVTILAGPTGAGKTH 486
YueF_Bs/1-369        ----------------------------------------------
F8FLY5_Pm/1-357      ----------------------------------------------
Upf0118_Ha/1-352     ----------------------------------------------
YtvI_Bs/1-371        ----------------------------------------------
Yct2_Bp/1-385        ----------------------------------------------
Q188A3_Cd/1-363      ----------------------------------------------

                       510       520       530       540       550
TqsA_Ec/1-344        ----------------------------------------------
YhhT_Ec/1-349        ----------------------------------------------
PerM_Ec/1-353        ----------------------------------------------
BARG_01524_Ba/1-658  LAEIWRSGTDALLVDPSNITEEAVNSAAERPVLIDNIGAEAFDETG 532
YueF_Bs/1-369        ----------------------------------------------
F8FLY5_Pm/1-357      ----------------------------------------------
Upf0118_Ha/1-352     ----------------------------------------------
YtvI_Bs/1-371        ----------------------------------------------
Yct2_Bp/1-385        ----------------------------------------------
Q188A3_Cd/1-363      ----------------------------------------------

                           560       570       580       590
TqsA_Ec/1-344        ----------------------------------------------
YhhT_Ec/1-349        ----------------------------------------------
PerM_Ec/1-353        ----------------------------------------------
BARG_01524_Ba/1-658  LFHLINSVRQHAAQGPGPSLLMTSRLWPANWNVKLPDLASRLKAAT 578
YueF_Bs/1-369        ----------------------------------------------
F8FLY5_Pm/1-357      ----------------------------------------------
Upf0118_Ha/1-352     ----------------------------------------------
YtvI_Bs/1-371        ----------------------------------------------
Yct2_Bp/1-385        ----------------------------------------------
Q188A3_Cd/1-363      ----------------------------------------------

                     600       610       620       630       640
TqsA_Ec/1-344        ----------------------------------------------
YhhT_Ec/1-349        ----------------------------------------------
PerM_Ec/1-353        ----------------------------------------------
BARG_01524_Ba/1-658  VVEIAEPDDMLLSGVIHKLFADRQVSVEPHVVSYLVSRMERSLLSA 624
YueF_Bs/1-369        ----------------------------------------------
F8FLY5_Pm/1-357      ----------------------------------------------
Upf0118_Ha/1-352     ----------------------------------------------
YtvI_Bs/1-371        ----------------------------------------------
Yct2_Bp/1-385        ----------------------------------------------
Q188A3_Cd/1-363      ----------------------------------------------

                         650       660       670
TqsA_Ec/1-344        ----------------------------------
YhhT_Ec/1-349        ----------------------------------
PerM_Ec/1-353        ----------------------------------
BARG_01524_Ba/1-658  IQIVDRLDRAALEQKSRITRALAAQILADMGQAG             658
YueF_Bs/1-369        ----------------------------------
F8FLY5_Pm/1-357      ----------------------------------
Upf0118_Ha/1-352     ----------------------------------
YtvI_Bs/1-371        ----------------------------------
Yct2_Bp/1-385        ----------------------------------
Q188A3_Cd/1-363      ----------------------------------
```

**Fig. S13 | Multiple sequence alignment of the AI-2 exporters.** Homologs considered include AI-2 exporters from *E. coli* (TqsA, YhhT and PerM), *Brucella abortus* (Uniprot: A0A7U8JIH7), *Paenibacillus mucilaginosus* (Uniprot: F8FLY5), *Bacillus subtilis* (YueF, Uniprot: O32095), *H.*

*andaensis* (Upf0118, Uniprot: A0A1W5X0D5), *B. subtilis* (YtvI, Uniprot O50485), *Bacillus pseudofirmus* (YCT2, Uniprot: Q04454) and *Clostridium difficile* (Uniprot: Q188A3).

**Table. S1| Primers used in the study**

| **Primers** | **Sequence (5'-3')** |
|---|---|
| TqsA_fwd | GAGGAATTAACCATGGCAAAGCCGATCATCACGC |
| TqsA_rev | GTGGCTCCAAGCGCTCTCTTTATTGAGATCGCTTAACAGAACGG |
| YdiK_fwd | GAGGAATTAACCATGGTAAATGTTCGTCAGCCCAGGG |
| YdiK_rev | GAGGAATTAACCATGGTAAATGTTCGTCAGCCCAGGG |
| YhhT_fwd | GAGGAATTAACCATGGAAACCCCTCAACCCGATAAAACG |
| YhhT_rev | GAGGAATTAACCATGGAAACCCCTCAACCCGATAAAACG |
| PerM_fwd | GAGGAATTAACCATGCTCGAAATGTTGATGCAATGGTATCG |
| PerM_rev | GTGGCTCCAAGCGCTTTCTTGCGCGATTTGCCCA |
